# *Pseudomonas aeruginosa* and *Candida albicans* both accumulate greater biomass in dual species biofilms under flow

**DOI:** 10.1101/2020.10.29.361139

**Authors:** Swetha Kasetty, Dallas L. Mould, Deborah A. Hogan, Carey D. Nadell

## Abstract

Spatially structured communities of microbes – biofilms – are widespread in nature, and biofilm-dwelling microbes often respond to their environments in ways that are different from their planktonic counterparts. Further, most natural biofilms are multi-species mixtures of microorganisms; the ecology of intra- and inter-species interactions in these consortia, and the resulting effects on total community properties, are often not well understood. A common site of polymicrobial biofilm infections is the lungs of patients with cystic fibrosis (CF). CF is a genetic disorder in humans that leads to colonization of the lungs by a variety of microorganisms, including *Pseudomonas aeruginosa* and *Candida albicans.* These opportunistic pathogens are frequently co-isolated from infected lungs, in addition to other infection sites including urinary and intravenous catheters. To study how these microbes behave together in biofilms, we developed a modified artificial sputum medium that is optically clear for use with microfluidic culture. In addition, we engineered strains with optimized fluorescent protein expression constructs allowing for single-cell resolution confocal microscopy. Using these tools and recently developed methods for spatial analysis of 3-D image data, we found that both *P. aeruginosa* and *C. albicans* display increased biovolume accumulation in multi-species biofilms relative to single-species biofilms. This pattern did not occur in planktonic co-culture and was thus specific to the biofilm environment. Interestingly, introduction of *P. aeruginosa* supernatants over dual-species biofilms strongly reduced *C. albicans* biovolume. This suggests that products that accumulate in batch culture were still inhibitory to *C. albicans* under a flow regime, but that they their *de novo* production in mixed species biofilms was not sufficient to inhibit *C. albicans* biofilm accumulation. Altogether our results indicate a critical impact of flow environment for the outcome of polymicrobial interactions and the need for high-resolution analysis of such communities in future work.

## Introduction

Microbial biofilm growth, even in mono-species contexts, involves the interplay of many biological and physical factors that are dynamic in space and time ^1,2,3^. In many natural environments, including numerous chronic infections, biofilms are multispecies mixtures whose collective properties may be difficult to predict from the those of each constituent’s mono-species biofilm growth. In the context of infection, the extent and kind of interactions among different biofilm-dwelling microbes also governs clinically relevant factors, such as drug resistance and virulence ^4^. For example, multispecies biofilm growth has been implicated in conjunctivitis^5^, tooth decay^6^, prosthesis and wound infections^7,8^, and respiratory diseases^9,10^. Clinical microbiologists are just starting to consider the multispecies nature of pathogenic biofilms, and its implications for prevention and treatment ^11^.

Exemplars of chronic, multi-species biofilm infections are those that occur consistently in the lungs of patients with cystic fibrosis (CF), a genetic disorder in humans as a result of mutations in the cystic fibrosis transmembrane conductance regulator. Disruption of this protein’s function results in pathologies throughout the body including the accumulation of highly viscous mucus in the lungs, which hinders normal mucociliary clearance. As a result, bacterial and fungal pathogens that would otherwise be easily removed from healthy lungs instead accumulate and lead to chronic infections ^12^. Chronic CF lung infections are caused by diverse and metabolically flexible multi-species consortia, and they are extremely recalcitrant to antibiotic and phagocytic clearance ^13^. While the ecology of the infecting species shapes the community and potentially has a profound influence on disease severity in the CF lung, it remains poorly understood ^9^. Given that the spatial interactions of pathogens can strongly affect disease outcome ^14^, we aimed to create an experimental model *in vitro* to investigate whether, and how, multi-species culture alters biofilm formation. Studies of multispecies biofilm formation and biofilm dynamics in general benefit tremendously from high resolution imaging, which allows for studying the cell-length-scale behaviors and higher order structures that contribute to the community’s cumulative growth, organization, and function. However, imaging live biofilms *in situ* is often difficult, if not impossible, in many natural contexts. A helpful strategy to mitigate this problem is to reconstitute key features of the *in situ* environment using an *in vitro* system that is more amenable to imaging.

As representatives of potentially interacting species in a polymicrobial CF infection, we chose here to study *Pseudomonas aeruginosa* and *Candida albicans,* both of which are commonly isolated from CF lung infections and believed to be important co-pathogens in patients ^15^. They are also thought to co-occur in other infection environments, including within trauma wounds and surrounding urinary catheters^16^. *C. albicans* is a polymorphic and opportunistic pathogen with the ability to form invasive hyphal filaments and drug resistant biofilms ^17^. *P. aeruginosa* is another opportunistic pathogen with diverse virulence mechanisms, to which biofilm formation contributes directly and indirectly ^18^. *P. aeruginosa-C. albicans* interactions are well studied in liquid and agar colony models. Among the primary findings from this literature, *P. aeruginosa* has been shown to inhibit the yeast-to-hyphal switch of *C. albicans* in liquid and agar colony cultures ^19^; *P. aeruginosa* has also been shown to preferentially attach to *C. albicans* hyphae in static culture, eventually killing them ^20^. Prior work has intimated a feedback loop whereby *C. albicans* produces ethanol, which increases biofilm formation, inhibits swarming motility, and enhances the production of antifungal phenazines on the part of *P. aeruginosa*. These phenotypes in turn cause downregulation of the central pathway that induces hyphal growth^21^ and inhibit mitochondrial activity, stimulating further ethanol production by *C. albicans*^22,23^. On the other hand, some *in vivo* experiments using a zebrafish model have indicated mutually enhanced virulence of the two species, suggesting that environmental shifts may have strong impacts on the properties of cocultures of these microbes^24^. Understanding how environmental shifts may change the nature of ecological interaction between *P. aeruginosa* and *C. albicans* is the primary goal of the present paper.

Using engineered strains with novel fluorescent protein constructs, and microfluidic culture with a modified synthetic sputum medium allowing for high-resolution imaging of *C. albicans* and *P. aeruginosa*, we show that their biofilm architecture and total biovolume accumulation differ strongly in monoculture versus coculture. Furthermore, surface-bound communities show qualitatively different dynamics from those in planktonic conditions. This result is robust to different clinical strains of *P. aeruginosa* and a variety of deletion mutants lacking factors known to participate in *P. aeruginosa-C. albicans* interactions. Furthermore, the interactions are highly dependent on spatial constraint and flow conditions in the milieu surrounding biofilm communities.

## Results

### Biofilm profiles in mono and dual culture

We aimed to characterize the architecture of mono-species and dual-species biofilms of *P. aeruginosa* and *C. albicans* under flow in a medium that represents the chemical composition of CF sputum. Synthetic cystic fibrosis media (SCFM), developed and refined by the Whiteley group ^25,26^, is a field standard for this purpose, but this medium is not optically clear due to the presence of reconstituted mucins. To generate an optically clear medium for imaging – and supported by data showing that *P. aeruginosa* does not degrade mucins itself ^27^ – we made a modified version of SCFM in which the major mucin glycans were substituted for mucin sugars; we term this modified medium artificial sputum media for imaging, or ASMi (see Materials and Methods). Each species’ growth profile was the same SCFM as it was in ASMi (SI Figure S1).

*P. aeruginosa* and *C. albicans* were modified by allelic exchange to contain a chromosomal construct for constitutive expression of *mKO-κ* (*P. aeruginosa*) or *mKate2 (C. albicans)* (See Materials and Methods). *mKO-κ* or *mKate2* were selected for these studies for their brightness and because they could be easily distinguished by fluorescence microscopy. The fluorescent protein expression constructs did not alter the growth rate of either species (SI Figure S2).

To investigate mono and dual culture biofilm growth under flow of ASMi, we inoculated derivatives of *P. aeruginosa* strain PA14 and *C. albicans* strains SC5314 either alone or together in microfluidic devices (see Materials and Methods). By visual inspection of confocal images, it was quickly clear that the architecture and total accumulation of both species were quite different in mono-versus dual-inoculated conditions (Figure 1). Monoculture biofilms of *C. albicans* contained scattered clusters of groups of elongated yeast, many pseudohyphae, and some true hyphae that spanned the height of the chamber (Figure 1B); in coculture however, *C. albicans* had largely formed true hyphae (Figure 1C), with higher biovolume density near the base of the biofilm (Figure 1D). Monoculture *P. aeruginosa* chambers contained small biofilms with compact microcolonies on the order of 10 μm in height (Figure 1D) and almost no detectable extracellular matrix carbohydrate (Pel) production relative to conditions that promote strong biofilm formation in this species (SI Figure S3). In coculture, *P. aeruginosa* biofilms localized to the hyphae of the highly filamentous *C. albicans* biofilms in proximity. The biovolume accumulation of *P. aeruginosa* in coculture appeared greater, particularly near the glass substratum, but also spanned the entire height of the chambers (Figure 1D).

**Figure 1.**
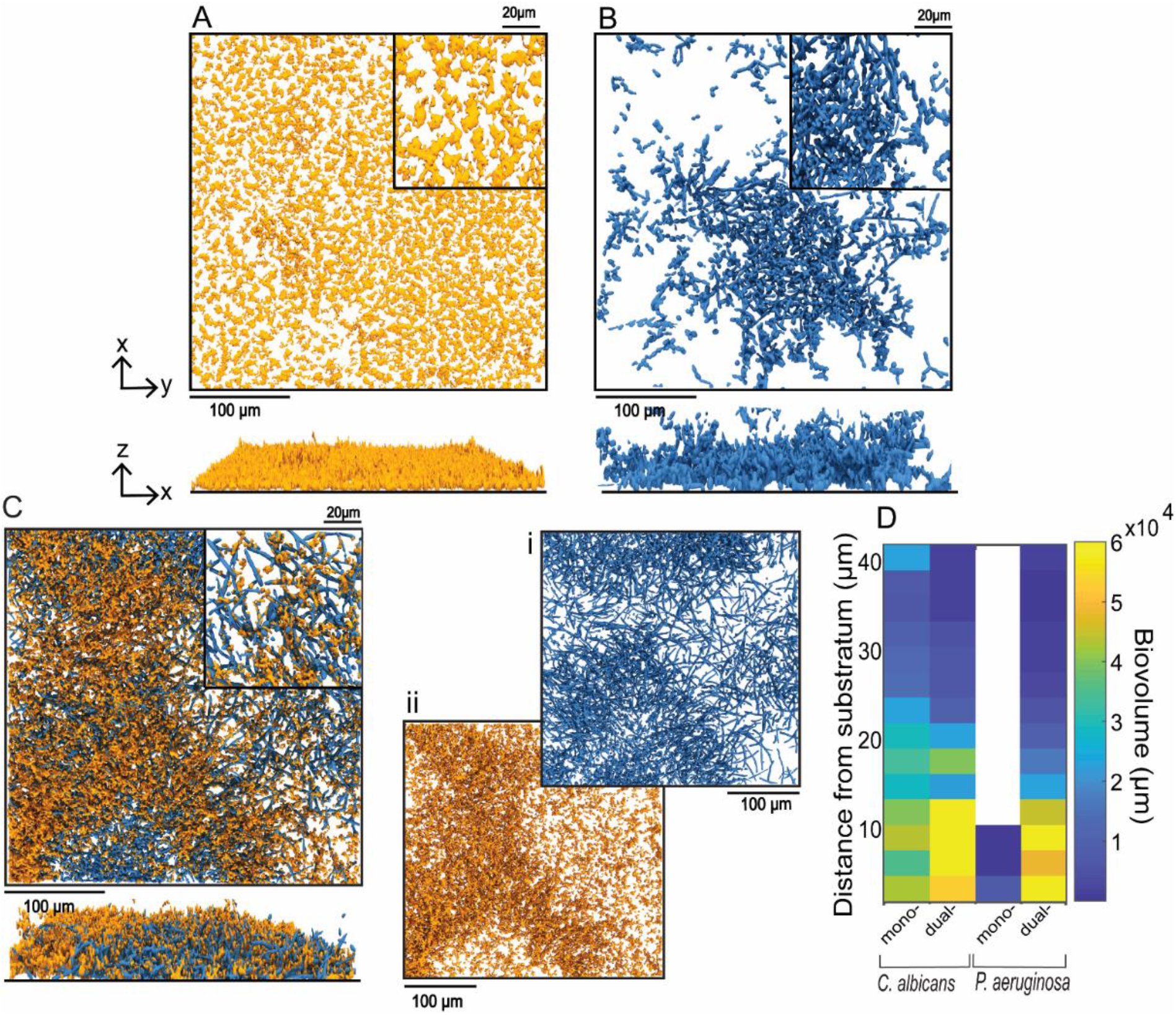
Representative images of mono- and dual-species biofilms of *P. aeruginosa* and *C. albicans.* 3-D renderings of 24-h old mono-species biofilms of (A) *P. aeruginosa* and (B) *C. albicans.* Bottom panels show side-views of the same images as those above them. (C) *P. aeruginosa-C. albicans* dual species biofilm at 24 hs i) Split channel of *C. albicans* biofilm from the *P. aeruginosa-C. albicans* dual species biofilm ii) Split channel of *P. aeruginosa* biofilm from the *P. aeruginosa-C. albicans* dual species biofilm. (D) Heat maps of the biovolume as a function of height for *P. aeruginosa* and *C. albicans* in mono- and dual-species biofilms from panels (A-C).

Image analysis confirmed that the total biovolume of both species increased substantially in coculture relative to monoculture (Figure 2A and 2B). Interestingly, in comparison experiments in which both organisms were cultivated in shaking liquid ASMi medium, the reverse pattern was seen for *C. albicans:* its population density was substantially lower in the presence of *P. aeruginosa* than in its absence (Figure 2C), recapitulating previously established antagonistic *C. albicans* – *P. aeruginosa* interaction in liquid growth conditions ^20,28,29^. The population density of *P. aeruginosa* did not change in the presence of *C. albicans* when in liquid culture (Figure 2D). We infer from this outcome that the increase in accumulation of both species in microfluidic coculture is specific to the biofilm environment. In order to verify that the increase in biovolume was indeed because of the presence of *P. aeruginosa*, we inoculated *C. albicans* on its own and grew it in microfluidic devices for 24 hs, then spiked *P. aeruginosa* into the chambers for one h, followed by a return to sterile ASMi medium. In control experiments, the same spiking procedure was performed, but with sterile ASMi medium. In the control treatment, *C. albicans* growth followed its normal monoculture profile, but in the experimental treatment, it increased after the introduction of *P. aeruginosa* (Figure 2E). We at first wondered whether the introduction of any mechanical disturbance might induce *C. albicans* to increase its biomass accumulation, but we saw no change in *C. albicans* biofilm architecture or biomass when inert fluorescent beads were introduced to the chambers instead of *P. aeruginosa* (SI Figure S4).

**Figure 2.**
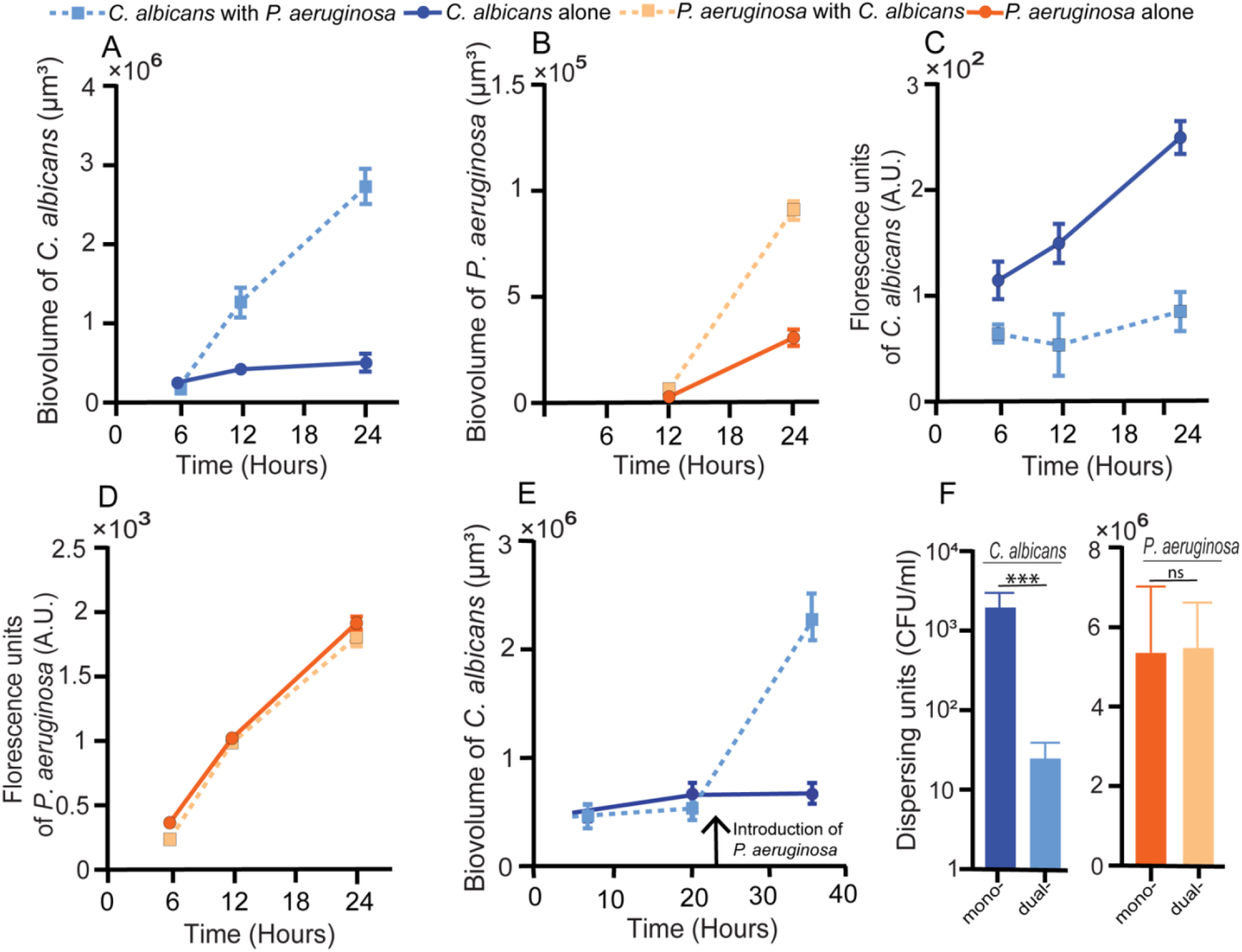
*P. aeruginosa and C. albicans* in mono- and dual-culture. (A) Biovolume of *C. albicans* in mono- and dual-species biofilms (n=24). (B) Biovolume of *P. aeruginosa* in mono- and dual-species biofilms (n=24). (C) Fluorescent counts of C. albicans in mono- and dual-species shaking liquid cultures (n=10).(D) Fluorescent counts of *P. aeruginosa* in mono- and dual-species shaking liquid cultures (n=10). (E) Biovolume of *C. albicans* biofilms initially grown in monoculture, with the addition of *P. aeruginosa* at the time point indicated by the vertical arrow. For the control, sterile media was added in place of *P. aeruginosa* (n =18). (F) Dispersing cells of *P. aeruginosa* and *C. albicans* in mono- and dual-species biofilms (p < 0.001; n=11). All error bars indicated are standard error.

Another potential reason for increased biovolume in coculture in addition to (but not mutually exclusive from) increased growth is higher retention of cells in the chambers, i.e. lower dispersal. To test whether dispersal of either species was reduced by the presence of the other, we collected media from the outlet of the microfluidic chambers and quantified the cells that were dispersing (see Materials and Methods)*. C. albicans* dispersal does indeed decrease in dual species biofilms (Figure 2F). Though it appears that *P. aeruginosa* dispersal stays the same in absolute terms (Figure 2F), it also decreased when the dispersal data are normalized to the amount of biovolume in the biofilm chamber (SI Figure S5).

### Fluid flow is critical for increased biofilm biomass of *P. aeruginosa* and *C. albicans* in dual culture

To try to determine the underlying mechanism involved in the *P. aeruginosa*-C*. albicans* biofilm interaction described in the previous section, we repeated the mono- and co-culture experiments above with mutants of *P. aeruginosa* that have been implicated in altered biofilm morphology or inter-species interaction in prior work. Analyses included mutants defective in the Pel exopolysaccharide production (Δ*pelA*^30,31^ and *ΔwspR^32^),* metabolic regulators and products important for biofilm formation (Δ*anr*^33^ and *Δphz^34^),* extracellular adhesins (Δ*bapA*^35^, *ΔpilY1^36^),* quorum sensing (Δ*lasR*^37^), and siderophore production *(ΔpvdApchE^38^).* All the *P. aeruginosa* mutants showed at least a modest increase in biofilm accumulation in the presence of *C. albicans* (Figure 3B), though for the weakest biofilm producers this result was not statistically significant. By contrast, *C. albicans* increased its accumulation by an order of magnitude or higher in biofilms with any of these mutants, maintaining the trend seen with wild type *P. aeruginosa* PA14 (Figure 3A, SI Figure S6). It is interesting to note that the amount of *P. aeruginosa* biofilm biomass present did not correlate with the degree of biomass increase in *C. albicans* (SI Figure S7); that is, any addition of *P. aeruginosa,* regardless of its native biofilm-producing capacity, was sufficient to produce a similar increase in accumulation of *C. albicans.* These results do not eliminate biofilm formation exoproducts or attachment factors as elements involved in *P. aeruginosa-C. albicans* interaction in co-culture, but they do suggest other important factors are at play.

**Figure 3.**
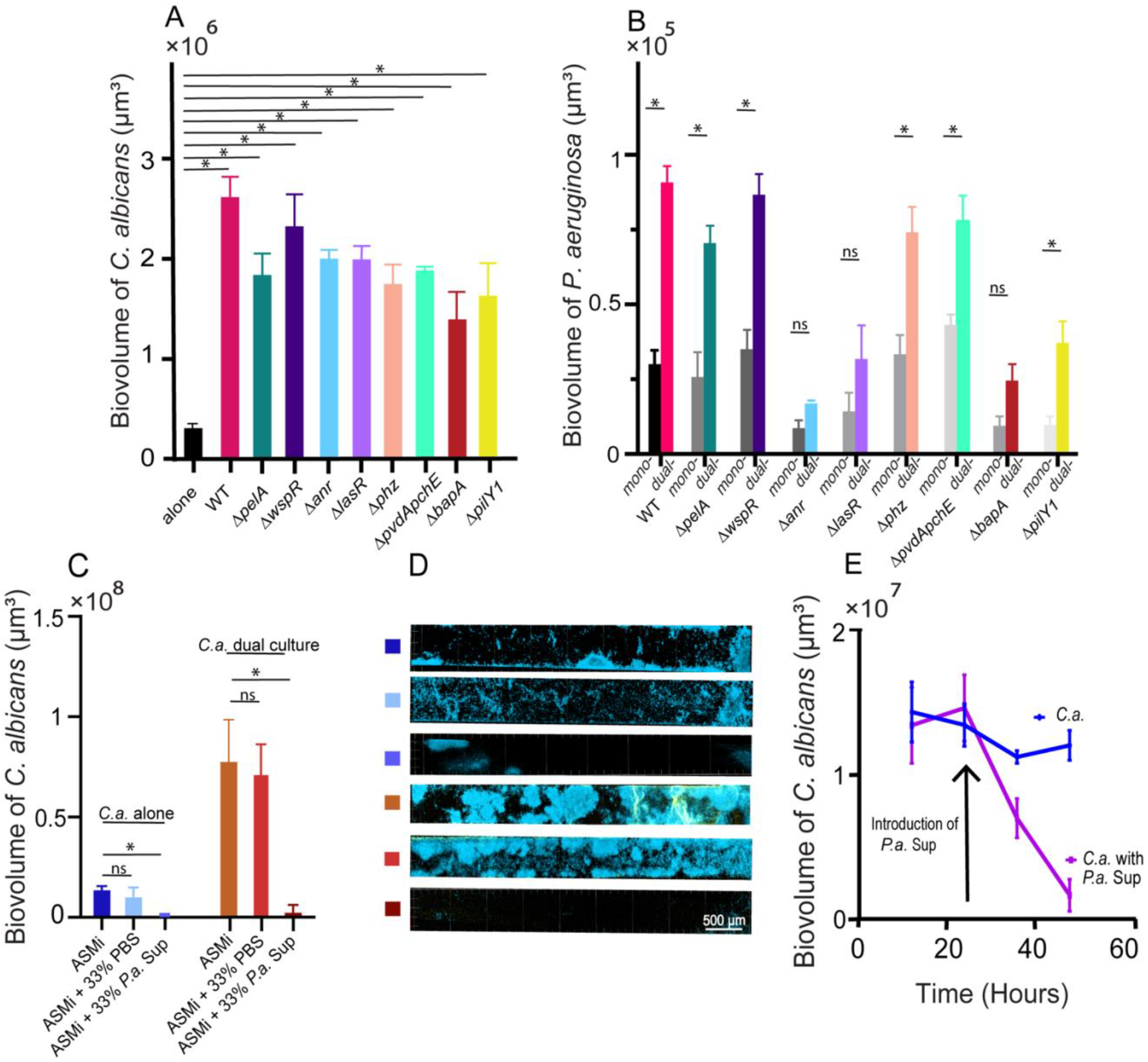
Deletion mutant assays and medium influent assays to explore the causes of mutual enhancement between *P. aeruginosa* and *C. albicans* in biofilms. (A) Biovolume of *C. albicans* grown in dual-species biofilms with *P. aeruginosa* deletion mutants at 24 hs (see main text for mutant descriptions, n=9-24). (B) Biovolumes of mono and dual species *P. aeruginosa* biofilms at 24 hs (n=9-24). (C) Biovolume of *C. albicans* mono species and dual species biofilms in the presence of different media influent. PBS is used as a control for addition of *P. aeruginosa* supernatants and media dilution. (n=6) (D) CLSM representative images of *C. albicans* and *C. albicans* + *P. aeruginosa* biofilms for conditions in C. (E) Biovolume of *C. albicans* biofilm initially grown in monoculture, followed by addition of *P. aeruginosa* supernatant at the time point marked by the arrow. For the control, sterile media was added in place of *P. aeruginosa* at the same time point (n=6). * denotes < 0.05. All error bars indicated are standard error.

As numerous studies have shown that secreted small molecule products from *P. aeruginosa* have an adverse effect on *C. albicans* growth and hyphae survival ^22,39^, we hypothesized that such secreted products were being removed from the system as a consequence of constant flow through the biofilm culture microfluidic devices. To test the role of exoproduct advection out of the biofilm chambers, we again grew *C. albicans* either alone or together with *P. aeruginosa*, with one of three different media treatments. In the first control treatment, chambers were run with ASMi for the entire experiment as described above. In a second control treatment, the media influent was replaced with a mixture of 2:1 ASMi:PBS. In the experimental treatment condition, the same procedure was performed, but the medium influent was replaced with a mixture of 2 parts ASMi to 1 part filter-purified supernatant from overnight liquid cultures of *P. aeruginosa* grown in ASMi. The goal of this experiment was to determine if reintroduction of *P. aeruginosa* secreted products into the biofilm chamber influent could reverse the biofilm biomass augmentation of *C. albicans* described above.

We found that the addition of *P. aeruginosa* supernatant caused a dramatic decrease in *C. albicans* biofilm accumulation, most pronouncedly when *C. albicans* was inoculated in coculture with *P. aeruginosa* (Figure 3C and 3D). Importantly, the PBS control treatment, in which *C. albicans* biomass was unaffected, demonstrates that addition of *P. aeruginosa* supernatants to the biofilm chamber influent does not reduce *C. albicans* accumulation due to a reduction of nutrient availability. Our first thought was that re-introduction of phenazines^40^ or rhamnolipid surfactants^41^ to the influent media might be primarily responsible for *C. albicans* biofilm biomass. However, supernatants from a *P. aeruginosa Δphz* mutant, which produces no phenazines, resulted in the same outcome as introduction of WT supernatants. Supernatants from *ΔlasR* mutant cultures, which lacks many quorum sensing regulated secreted products including phenazines and rhamnolipids, were partially effective compared to the effects of WT supernatants, suggesting that the effects were likely multifactorial (SI Figure S8). The observation that *P. aeruginosa* supernatant remained partially effective after protease treatment supports the model that multiple factors contributed to the effects on biovolume accumulation. Interestingly, it is worth noting that all supernatant activity was lost after boiling.

Previous studies have suggested that *P. aeruginosa* supernatants alone are not sufficient to inhibit *C. albicans* viability, but did not specifically examine their effect on biomass retention to surfaces for biofilm production ^20,29^. To explore this possibility we grew biofilms of *C. albicans* in monoculture for 24 hs under regular ASMi flow, allowing them to reach their typical biomass steady state in biofilm chambers; after this 24 h period the medium influent was switched to a 2:1 mixture of ASMi and *P. aeruginosa* culture supernatant. After introduction of the supernatant, the *C. albicans* biofilms decreased linearly in total biomass nearly to zero in 24 h (Figure 3E). Altogether these results indicate that the increased biomass accumulation of *C. albicans* in the presence of *P. aeruginosa* is dependent on the mass transport of *P. aeruginosa* secreted products out of the system. When these supernatants are reintroduced, of *C. albicans* accumulation in the chambers ceased, and any pre-existing biomass was greatly reduced over time. This could be due to inhibited growth of *C. albicans*, inhibited attachment of *C. albicans* to substrata, induced dispersal, or a combination of these factors.

### *P. aeruginosa – C. albicans* interaction is robust to CF isolate variation

After documenting that PA14 wild type could induce an increase in biofilm biomass accumulation of *C. albicans,* we were curious to see whether this effect was consistent across recent CF clinical isolates of *P. aeruginosa* as well. To explore this question, we obtained *P. aeruginosa* clinical isolates from a patient who was infected with both *P. aeruginosa* and *C. albicans,* and we grew them in mono- or coculture with *C. albicans* in our microfluidic model under flow of ASMi. We found that *C. albicans* biofilm increased significantly in coculture with all clinical isolates, consistent with the results reported above for PA14 wild type (Figure 4A). Likewise, for all but one isolate, the biofilm growth of *P. aeruginosa* was greater in coculture with *C. albicans* than it was in monoculture (Figure 4B).

**Figure 4.**
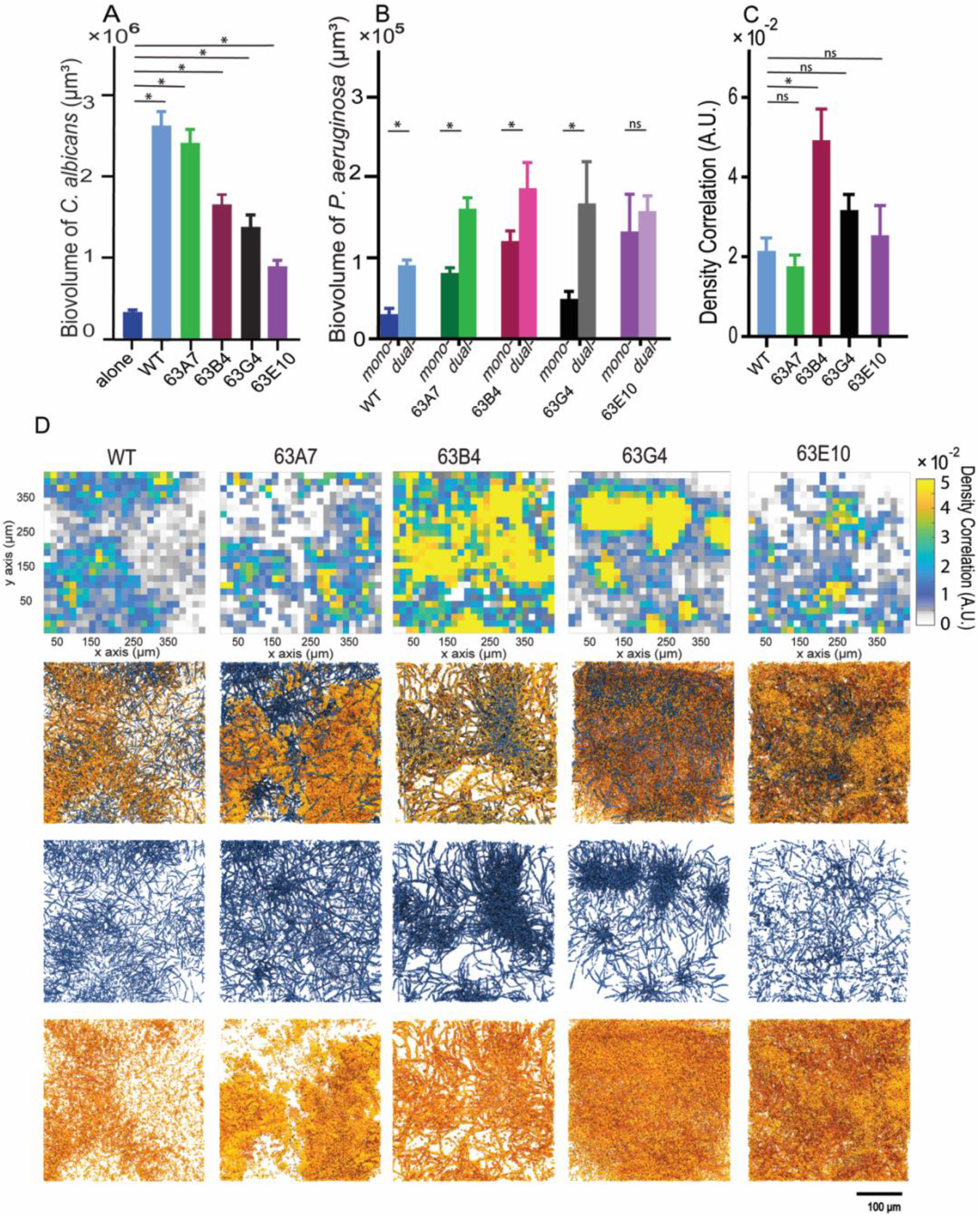
Biomass accumulation, density correlation analysis, and visualization of *C. albicans* in co-culture with different CF clinical isolates of *P. aeruginosa.* (A) Biovolume of *C. albicans* grown as dual species biofilms with *P. aeruginosa* clinical isolates along with PA14 wildtype for comparison at 24 hs. (n=18). (B) Biovolumes *P. aeruginosa* clinical isolates in monoculture and dual culture with *C. albicans* at 24 hs (n=18). (C) Global density correlation measurements of *P. aeruginosa* WT and clinical isolates and *C. albicans* biofilms (n=6). * denotes p < 0.05. (D) Visualization of dual-species biofilms of *P. aeruginosa* and *C. albicans;* from top to bottom: spatially resolved density correlation; 3-D renderings of dual species biofilms; *C. albicans* channel split; *P. aeruginosa* channel split.

Though all clinical *P. aeruginosa* isolates prompted an increase in *C.* albicans biofilm accumulation, there was some variance in the degree to which this was the case (Figure 4A). This variation made us wonder if the spatial association between *C. albicans* and different clinical isolates of *P. aeruginosa* might differ as well. To assess this possibility, we grew the different clinical isolates together with *C. albicans*, acquired high resolution images of coculture biofilms, and quantified the spatial co-occurrence of the two species via their density correlation^42^. When averaged across all image replicates, the spatial correlations between *C. albicans* and clinical isolates of *P. aeruginosa* generally were not different from that between *C. albicans* and PA14 wild type (Figure 4C). After visualizing the density correlation measurement at high spatial resolution, on the other hand (Figure 4D), it was clear that for some clinical *P. aeruginosa* isolates, the spatial association with *C. albicans* was homogenous, while for others it was patchy. The significance of this result for the infection ecology of these two species is not yet clear, but it is notable that among isolates of *P. aeruginosa* from the same patient, the architecture of joint biofilms with *C. albicans* can differ substantially at the micrometer scale (Figure 4D) even when they appear to be the same or similar when averaged on a larger spatial scale (Figure 4C).

## Discussion

Interest in multi-species biofilms including microbes from different domains of life has been intensifying in recent years, as it is increasingly appreciated that many microbial communities – both inside and outside of host organisms – are polymicrobial^43^. One of the most highly referenced examples of polymicrobial infections are those within the lungs of patients with CF, and two of the common members of these communities are the opportunistic pathogens *P. aeruginosa* and *C. albicans*^12^. Here we sought to compare the kinetics of biovolume accumulation in mono- and dual-species biofilms of these two organisms using a new model of biofilm growth under flow of optically clear artificial sputum medium. We demonstrated a marked increase of biofilm biomass accumulation as well as a decrease in dispersal of both species in dual-culture relative to monoculture. These results were robust to a variety of mutant and clinical strain backgrounds of *P. aeruginosa*, and they contrast with the findings of some previous studies of these two organisms in static liquid or agar colony culture ^44,45^. We identify an important element driving the increase in biomass accumulation as fluid flow in the dual-species biofilm milieu, which is a key novelty of this experimental approach for the study of *P. aeruginosa-C. albicans* interactions.

Extensive prior work has shown that *P. aeruginosa* and *C. albicans* interact with each other through a complex web of secreted factors, including phenazines, siderophores, ethanol, and quorum-sensing autoinducers, which altogether alter environmental iron availability, pH, and oxygen tension (Figure 5A). Under static culture conditions (i.e. liquid batch culture or agar colonies), the net result of these interactions is usually antagonism of *P. aeruginosa* against *C. albicans.* It is important to note as well that secreted factors from each species have different and sometimes opposite effects on each other’s propensity to produce biofilms or to remain in a dispersive, planktonic state ^28,40^. As noted above, when flow – known to impact microbial physiology and surface interaction – is introduced into the two-species system, we see increased filamentation of *C. albicans* and increased biofilm biomass accumulation by both species, accompanied by a decrease in dispersal.

**Figure 5.**
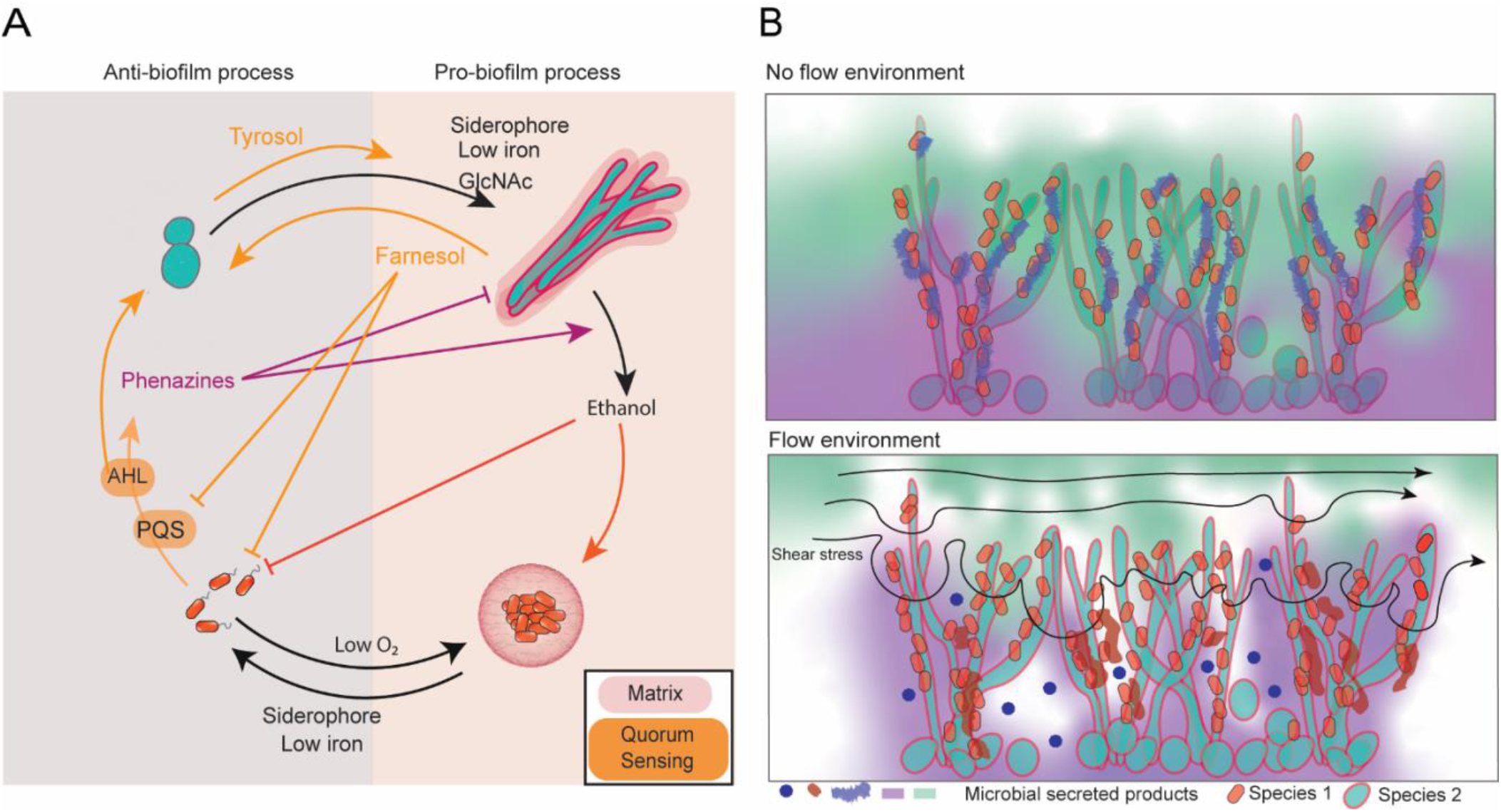
Interspecies interactions and effect of flow on biofilm properties. (**A**) Documented effects of extracellular molecules effecting *P. aeruginosa* and *C. albicans.* (**B**) Representation of potential effects on flow including changes in the secreted solute environment, changes in secreted products and production of new extracellular chemicals. Flow patterns can also change in different regions resulting in difference in chemical concentrations and shear stress.

While at first glance this may give the impression of mutual benefit, it is also possible that the two species are simply competing for access to space and resources by upregulating adhesion factors^46–48^. But why is *P. aeruginosa* no longer able to directly antagonize and kill *C. albicans*, as has been shown previously in static culture? We speculate that introduction of flow fundamentally changes the secreted solute environment created by the two organisms, perhaps with some secreted factors more strongly retained in the biofilm matrix than others, and that this change in solute environment relative to static culture shifts the ecological pattern of biomass accumulation to one in which both species are augmented. It is also possible that over time the dual-species biofilms become densely packed enough to block flow within some regions^49,50^, allowing secreted products and variation in iron/oxygen availability to accumulate in a patchy manner that contributes to induction of biofilm production by both species (Figure 5B). The precise spatial patterns of exoproduct accumulation in relation to cells and the highly complex matrix that *Candida* secretes is an important area for future work^51–53^.

Our deletion mutant analysis included all of the major classes of behavior in *P. aeruginosa* currently known to mediate solute-based interactions with *C. albicans*, but in all cases the presence of *P. aeruginosa* caused qualitatively the same increase in *C. albicans* biofilm. This suggests that there may be other factors in addition to flow-mediated changes in solute environment contributing to our results. For example, the introduction of shear stress under flow is an entirely new environmental stimulus relative to static culture, and one which is known via extensive work to be crucial to microbial ecology and evolution^54–59^. Flow regime can dramatically alter the morphology and resilience of bacterial biofilms down to their cellular resolution architecture^60,61^, with important implications for pathogenesis in the case of infections^62^. Adaptation to the challenges of flow at sub-millimeter spatial scales has influenced the evolution of bacterial surface motility^2^, optimal growth rate in porous media^63^, surface colonization mechanisms^64–66^, extracellular matrix secretion^67,68^, bacterial cell shape^65,69–71^, planktonic aggregate formation^72^, and biofilm community assembly and function^63,73–76^, to name just some examples among many.

In light of our results it is important to note that flow regime has documented effects on biofilm formation for both *P. aeruginosa* and *C. albicans*. The surface residence time of *P. aeruginosa*, for example, increases linearly as shear stress increases^77^, and flow promotes upstream surface motility in addition to the formation of biofilm aggregates ^78^. *P. aeruginosa* has also recently been shown to be highly responsive to mechanical stress induced by flow, with downstream effects on biofilm formation that have yet to be fully clarified^36,79^. There has been less investigation of the effects of shear flow on *C. albicans* biofilms: existing work does not agree completely on whether shear stress increases total biomass of *C. albicans* biofilms but does agree that biofilms formed under shear are more highly compacted and physically robust relative to those grown in static conditions^80^. It is also feasible that by de-emphasizing the impact of diffusible solutes, dual-culture biofilms under flow reveal previously underappreciated effects of direct contact-mediated interactions between *P. aeruginosa* and *C. albicans*, which we document here to be in tight association with each other. Importantly, given that dual-species culture produced substantial biomass accumulation for both species relative to monoculture under the same flow conditions, flow-induced shear cannot on its own explain our results. Rather we infer that a combination of physical forces resulting from flow in addition to biological interaction between the two species must be responsible for the results obtained here. Dissecting the precise molecular mechanisms of these inter-species interactions is an important area for future study, that may bear directly on the outcome of multi-species biofilm growth in the context of infection.

Beyond their prevalence in lung infections among patients with CF, *P. aeruginosa* and *C. albicans* individually are among the most common agents of nosocomial infection currently known^16^. They are both frequently isolated from device-related infections including implanted medical devices, prosthetic implants in wounds and joint replacements, and urinary catheters ^16^. Both species participate in multi-species infections, for example, with *Staphylococcus* spp.^81–83^, with *Streptococcus* spp.^84,85^, and with each other^39^. Reports of dual isolation of *P. aeruginosa* and *C. albicans* are increasingly reported in the clinical literature in sites such as ventilator tubing ^86^, and our results of biofilm dual-culture in microfluidic devices suggest that dual *Pseudomonas*/*Candida* biofilms may be especially problematic in this setting because they tend to accumulate more biofilm biomass together than alone. Such rapidly accumulating biofilms can potentially clog catheter flow environments and seed systemic infections as cells disperse from the device-attached biofilm into the bloodstream.

Though recent studies have made tremendous strides in imaging microbiomes within *in situ* samples that have been fixed ^87–90^, dissecting live microbial community structure in space and time within native environments remains a challenging task and one of the important frontiers of modern microbiology. Here we use an *in vitro* model with medium tuned to the CF sputum environment to assess live biofilm population dynamics and find that this step toward environmental realism has a strong impact on the ecology of dual-species biofilms of *P. aeruginosa* and *C. albicans.* Many native factors are still missing, however: the mucosal environment is quite different in the native lung, for example, and recent work has suggested that mucus has a strong impact on *P. aeruginosa* physiology, including reducing its propensity toward virulence and biofilm formation ^91,92^. Though not an exact match to the *in situ* infection environment, our system nevertheless suggests that modest changes to the environmental context in which multispecies interactions are studied can have a large impact on the observed outcome, namely in this case, a shift toward far higher accumulation of biofilm on the part of *P. aeruginosa* and *C. albicans* when they are together versus when they are alone. On the basis of this observation we speculate that pushing toward realism and high-resolution image analysis of biofilm communities will yield important and unexpected insights for many other microbial systems of interest.

## Materials and Methods

### Strains and media

Supplementary Table S1 includes a full strain and plasmid list for this study. Strains of *P. aeruginosa* are either derivatives of strain PA14 or clinical isolates. Strains of *C. albicans* are derivative of CAI4. All strains were grown on LB (10 g tryptone 5g NaCl 5g yeast extract per liter) and ASMi (*P. aeruginosa*) or YPD (10 g yeast extract 20 g peptone and 20 g dextrose per liter) and ASMi (*C. albicans*). The media recipes and concentrations of reagents used for ASMi are listed in the Supplementary Information. All chemicals and reagents were purchased from Millipore Sigma unless otherwise stated.

### Plasmid and strain construction

All restriction enzymes and ligase were purchased from New England Biolabs, and PCR reagents were purchased from BioRad. The *P. aeruginosa* tandem codon-optimized version of *mKO*-κ was custom synthesized by Invitrogen. The construct contains two copies of *mKO-κ* in tandem, each with its own ribosome binding site, and with different codon composition to prevent excision by recombination. Florescent *P. aeruginosa* derivatives were constructed by amplification of the flanking regions upstream and downstream of the Tn7 *att* site and fusion of the custom fluorescent protein construct to a synthetic *tac* promoter for high expression from a single chromosomal locus. This fused construct was cloned into the pMQ30 plasmid used for allelic exchange in *P. aeruginosa*^93^. This plasmid was then introduced into *Escherichia coli* S17-λpir by electroporation, conjugated into *P. aeruginosa,* and recombinants were obtained using selection on gentamicin and sucrose counter-selection for loss of the integrated plasmid backbone. For *C. albicans*, a single codon-optimized version of *mKate2* was custom-synthesized by Invitrogen. The RP10 integrative plasmid, pACT-GFP^94^, has been shown to have constant expression levels through *C. albicans* growth cycle. We replaced the GFP in pACT-GFP^94^ with *mKate2.* For transformation into *C. albicans*, the *mKate2-* containing plasmid was linearized by BglII restriction digest, concentrated using the Zymo Research DNA Clean & Concentrator-5 kit (Cat. # 11-303), and 1 μg was electroporated into electrocompetent *C. albicans* CAI4 prepared as previously described^95^. Prototrophic recombinants were selected for on uracil drop-out media.

### Liquid growth curve and florescence measurements

*P. aeruginosa* strains were grown at 37°C shaking in LB overnight prior to growth curve experiments. The following morning cultures were back-diluted to an OD^600^ of 0.01 in ASMi in 10mL glass tubes with 2mL media (for fluorescence growth curves) or 50mL Falcon tubes with 30mL media (optical density growth curves), rotating at 250 rpm on an incubated orbital shaker at 37°C. *C. albicans* strains were grown at 30°C shaking in YPD overnight prior to growth curve experiments. They were cultivated overnight at 30°C to maintain cells in yeast form prior to the start of growth curve or biofilm experiments (see below). The following morning cultures were back-diluted to an OD^600^ of 0.01 in ASMi in 10mL glass tubes with 2mL media (for fluorescence growth curves) or 50mL Falcon tubes with 30mL media (optical density growth curves), rotating at 250 rpm on an incubated orbital shaker at 37°C. Fluorescence measurements were made using a Synergy Neo2 every 6 hs. A 543-nm excitation source was used to excite *mKO-κ*, and a 594-nm excitation source was used to excite *mKate2.* Optical density measurements were made every h using a benchtop spectrophotometer (CWA Biowave CO8000 Cell Density Meter).

### Microfluidic device assembly

The microfluidic devices were made by bonding polydimethylsiloxane (PDMS) chamber molds to size #1.5 cover glass slips (60mm X 36mm [LxW], Thermo-Fisher, Waltham MA) using standard soft lithography techniques^96^. Each PDMS mold contained 4-5 chambers, each of which measured 3000μm x 500μm x 75μm (LxWxD). To establish flow in these chambers, media was loaded into 1mL BD plastic syringes with 25-gauge needles. These syringes were joined to #30 Cole-Parmer PTFE tubing (inner diameter 0.3 mm), which was connected to pre-bored holes in the microfluidic device. Tubing was also placed on the opposite end of the chamber to direct the effluent to a waste container. Syringes were mounted to syringe pumps (Pico Plus Elite, Harvard Apparatus), and flow was maintained at 0.1 μL per min for all experiments.

### Biofilm growth, matrix staining and CFU counts

Overnight cultures of *P. aeruginosa* were grown at 37°C shaking in LB, and overnight cultures of *C. albicans* were grown at 30°C shacking in YPD prior to the start of biofilm experiments. Cultures of both strains were normalized to an OD600 of 0.05 in ASMi media. If dual culture biofilms were to be started, equal volumes of OD-equalized strains were mixed and inoculated into a microfluidic chamber (completely filling its inner volume), and then allowed to rest for 1 h at 37°C to permit cells to attach to the glass surface. The devices were then run at 0.1 μL per min at 37°C, and imaged by confocal microscopy (see below) at time intervals that varied per experiment as noted in the main text. All experiments were repeated with at least 5 biological replicates with 3 or more technical replicates on different days. Total replicates for each experiment are noted in the figure legends for each data set in the text and Supplementary Information.

Wisteria floribunda lectin stain (Vector Labs®) conjugated to fluorescein dye was used to visualize pel polysaccharide produced by *P. aeruginosa^31^.* The lectin was added to the media in syringes for these experiments such that biofilms would be exposed to the lectin -dye conjugate for the entire period of biofilm growth (20 μL stock lectin solution per mL of media, per manufacturer’s protocol recommendation from a stock solution of 2mg/mL dye conjugate). Biofilms were inoculated as noted above for these experiments and grown for 24 hs prior to imaging.

To compare growth rates of *P. aeruginosa* and *C. albicans* in turbid SCFM and newly optimized ASMi, both species were grown overnight, *P. aeruginosa* in LB at 37°C and *C. albicans* in YPD at 30°C in 10mL glass tubes with 2mL media. The following morning cultures were back diluted to an OD^600^ of 0.01 in either SCFM or ASMi in 50mL Falcon tubes with 30mL media, rotating at 250 rpm in an orbital shaker at 37°C. 1mL of culture was taken from the Falcon tube at different time points, serial dilution was performed and plated on LB agar for *P. aeruginosa* and YPD agar for *C. albicans*. The number of colony forming units from each plate was recorded and used to calculate growth rates measured by CFU/mL per time.

To measure dispersed cell counts, biofilms of both species were grown as noted above in ASMi media for 24 hs, after which the outlet tubing of the microfluidic device was changed to ensure we were measuring dispersal only from the biofilms within the chambers themselves. The flow rate was increased to 500 μl per min and outflow collected. Serial dilutions were performed and plated on LB agar for *P. aeruginosa* and YPD containing 50 μg / ml chloramphenicol for *C. albicans.* The number of colony forming units (CFUs) from each plate was recorded and used to calculate the CFU/mL culture density emerging from the chambers. This experiment was repeated for 11 biological replicates with independent overnight cultures.

### Supernatant experiments

*P. aeruginosa* wild type or mutants were grown overnight in 10mL glass tubes with 2mL of ASMi at 37°C and 250 rpm on an incubated orbital shaker. The overnight cultures were centrifuged at 10000 rpm for 1 min. The supernatant was collected and passed through a 0.2 μm filter. For treatment with Proteinase K and DNAse I (New England Biolabs®), the respective enzyme was added to filtered supernatant according to the NEB® protocol and allowed to incubate for 1 h. For the boil treatment, the filtered supernatant was boiled at 100°C for 15 mins. Supernatants were then added to fresh sterile ASMi at a ratio of 1:2. The mixed media was then used to grow *C. albicans* biofilms in microfluidic chambers, or in a separate experiment, introduced to 24 h old *C. albicans* biofilms (see main text). In both cases biofilms were inoculated and cultivated as described above.

### Microscopy and image analysis

Biofilms inside microfluidic chambers were imaged using a Zeiss LSM 880 confocal microscope with a 40x/1.2NA or 10x/0.4NA water objective. A 543-nm laser line was used to excite mKO-κ, and a 594-nm laser line was used to excite mKate2. A 458-nm laser line was used to excite Wisteria floribunda lectin stain in the case of Pel quantification experiments. All quantitative analysis of microscopy data was performed using BiofilmQ ^42^. 3-D renderings of biofilms in Figure 1 and Figure 4 were made using Paraview.

### Statistics

All statistical analyses were performed in GraphPad Prism. All reported pairwise comparisons were performed using Wilcoxon signed-rank tests, and multiple comparisons were performed by Wilcoxon signed-rank tests with Bonferroni correction. All error bars indicated are standard error.

## Acknowledgements

CDN and DAH conceived and supervised the project. SK, DLM, DAH, and CDN designed experiments. SK performed experiments and image analysis. SK and CDN finalized figures. DLM and DAH contributed critical reagents. SK, DLM, DAH, and CDN wrote the paper.

## Supplementary Information

### Supplementary Figures

**SI Figure S1.**
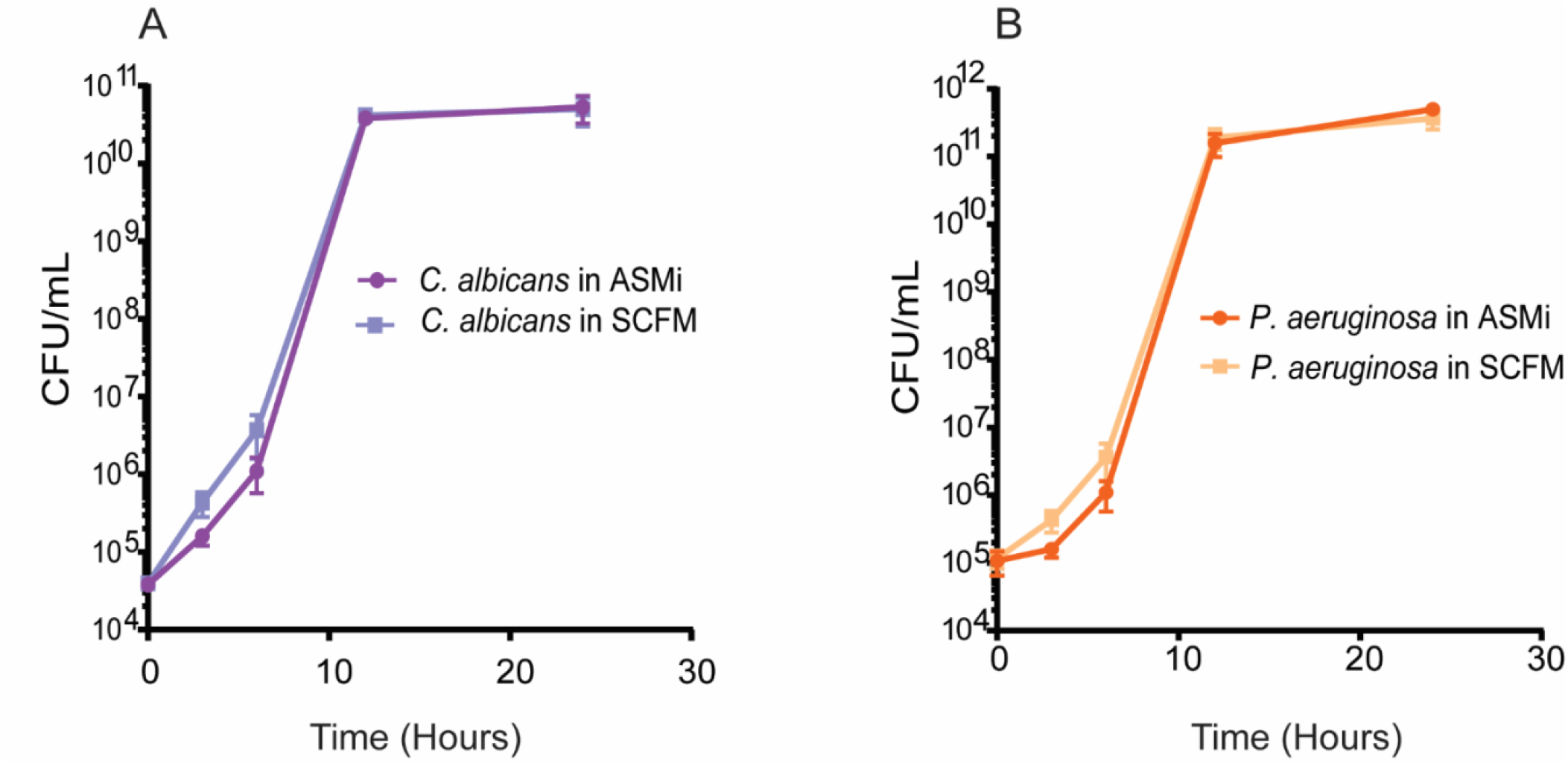
Growth curves of (A) *C. albicans* and (B) *P. aeruginosa* in standard SCFM medium with reconstituted mucin and in ASMi medium containing only mucin glycans without full-length mucin polymers (n = 6). All error bars indicated are standard error.

**SI Figure S2.**
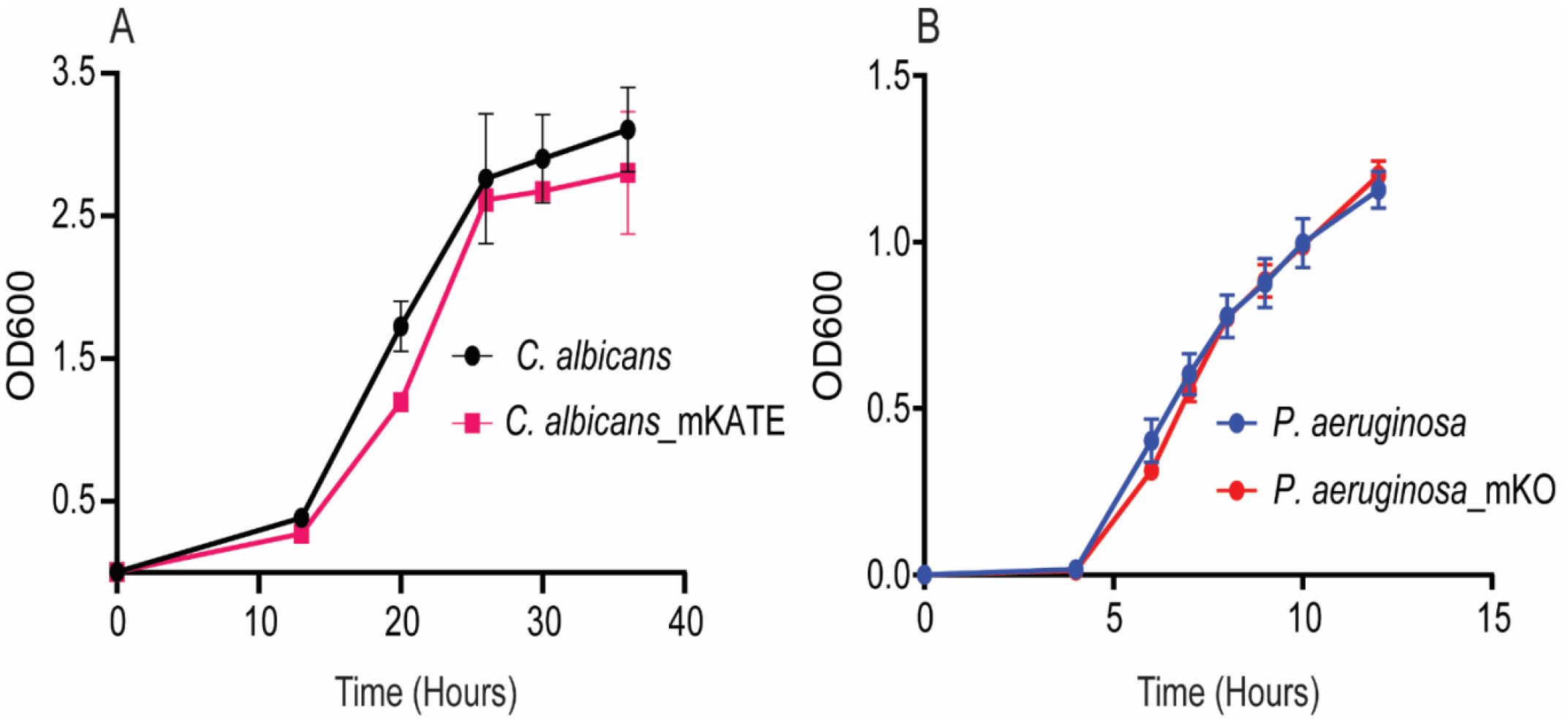
Growth curves in ASMi media of wild type (A) *C. albicans* SC5314 wild type or *C. albicans mKate2* and (B) *P. aeruginosa* PA14 wild type or *P. aeruginosa _mKO-κ* (n=6). All error bars indicated are standard error.

**SI Figure S3.**
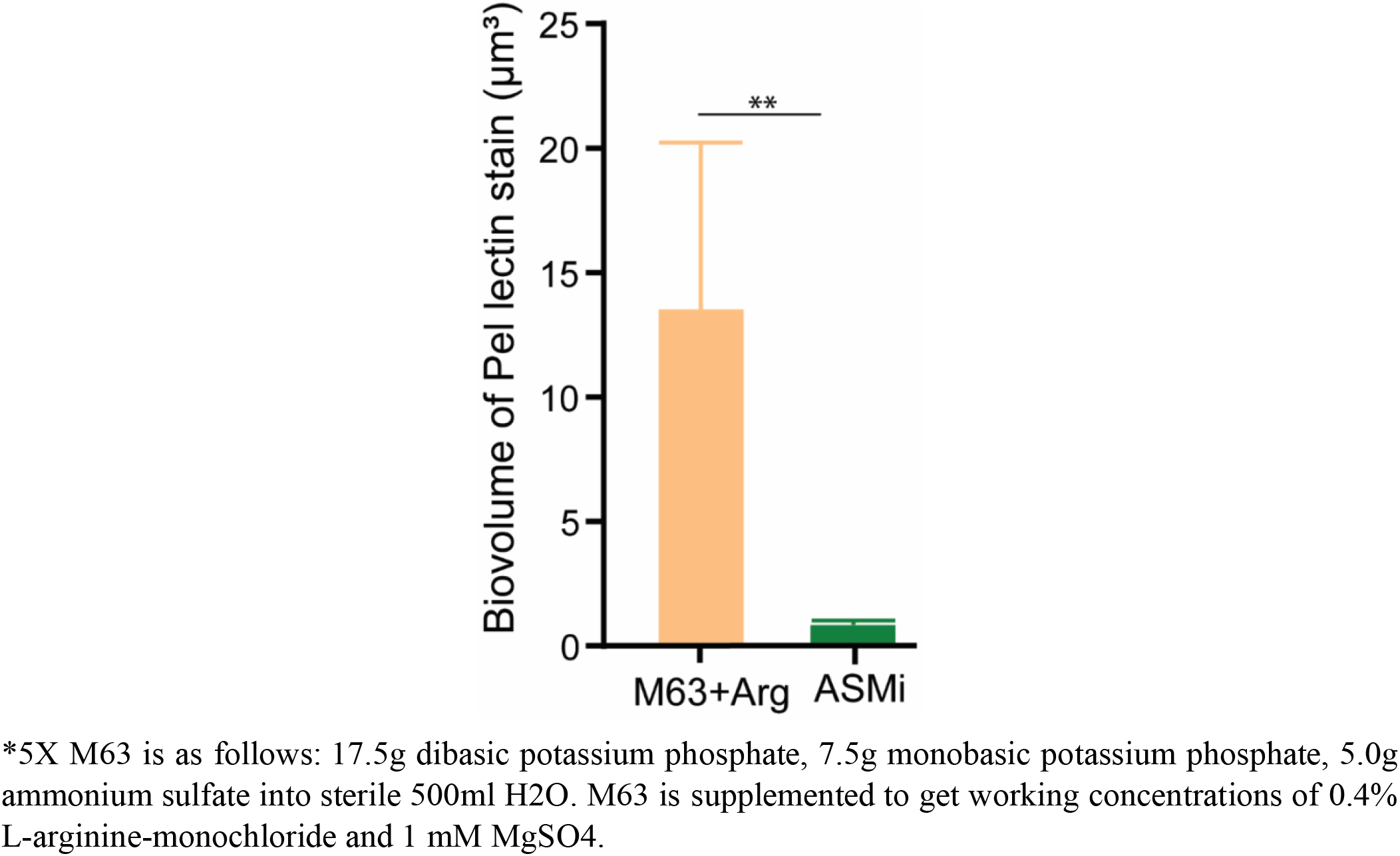
The matrix polysaccharide Pel was stained and quantified in *P. aeruginosa* strain PA14 biofilms using florescence marker bound to Pel-specific lectin in M63 media plus arginine, a biofilm assay media ^97^) and ASMi medium (p<0.001; n=6). Reported pairwise comparisons determined by Wilcoxon signed-rank test. All error bars indicated are standard error.

**SI Figure S4.**
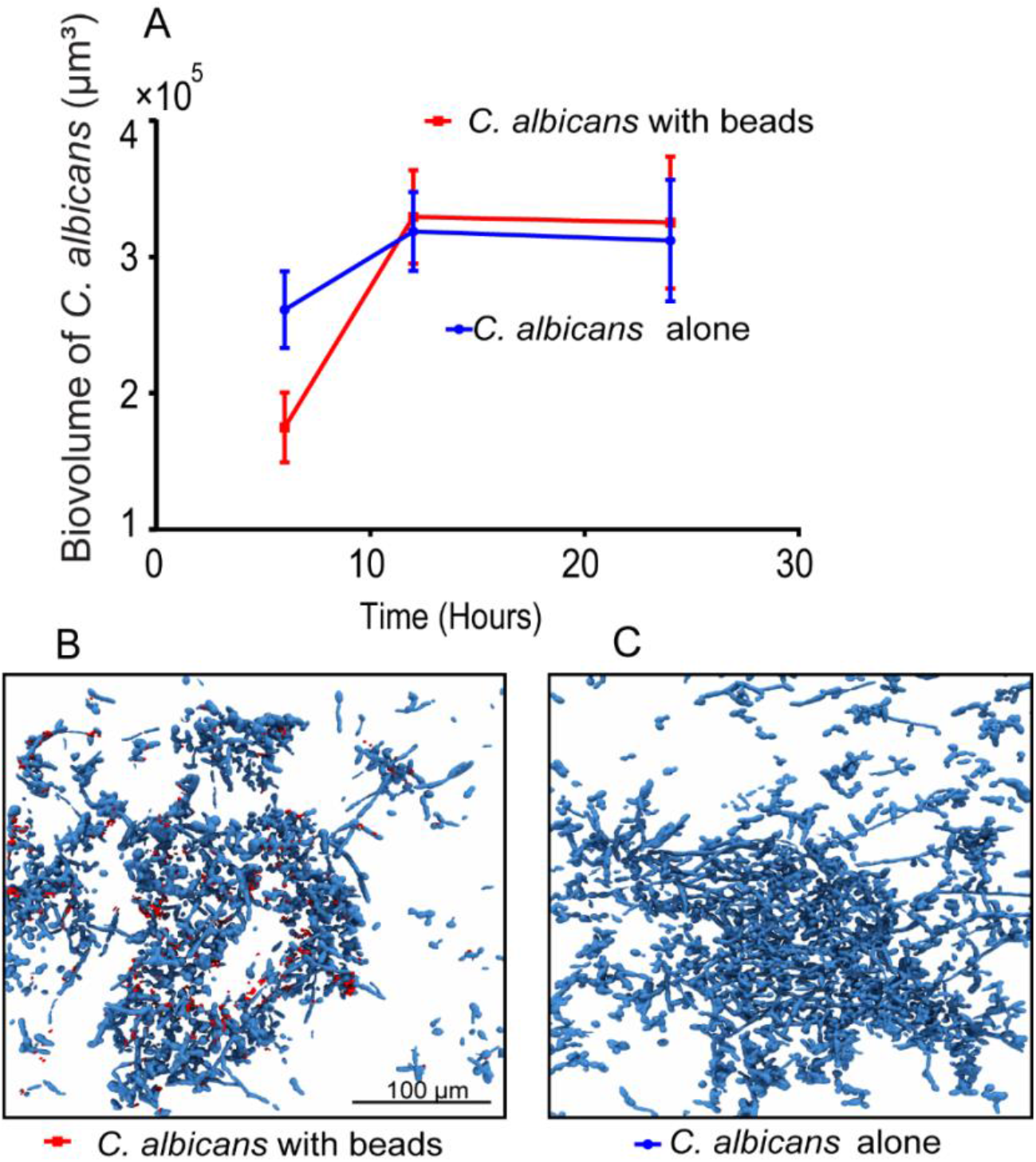
Inert fluorescent beads were introduced to *C. albicans* expressing *mKate2* biofilms to test for the effect of mechanical disturbance on *C. albicans* biofilm growth. (A) Biovolume of *C. albicans* with and without fluorescently labeled beads added to the influent media (n=6). (B) Representative image of *C. albicans* biofilms grown with fluorescently labeled beads (shown in red) at 24 h time point (C) Representative image of *C. albicans* biofilm in the absence of fluorescent beads in influent media at 24 h time point. All error bars indicated are standard error.

**SI Figure S5:**
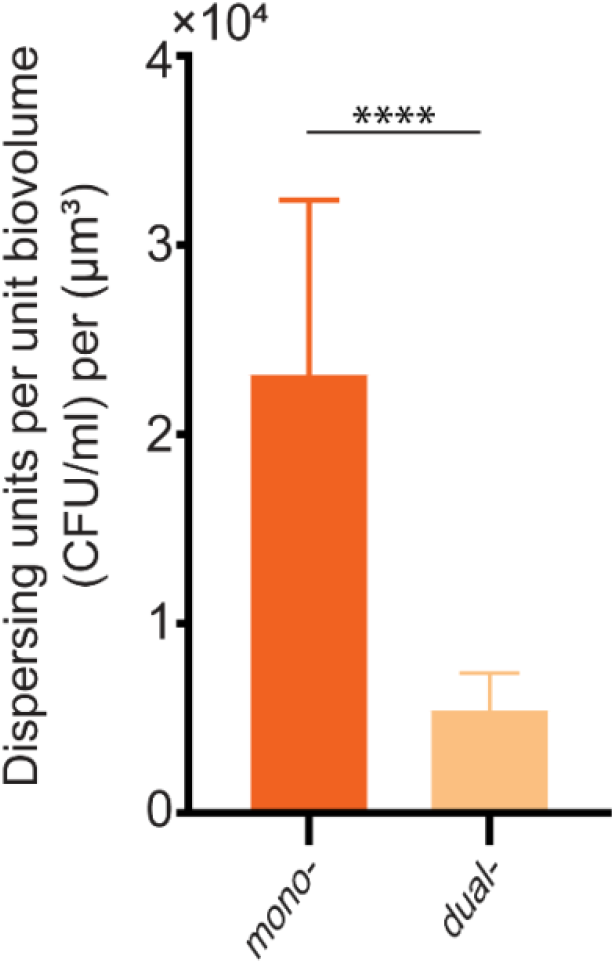
Dispersing *P. aeruginosa* CFUs units per unit biovolume. Dispersing units of *P. aeruginosa* obtained from plating cells from the outflow of the microfluidic chamber normalized to biovolume of cells present in the microfluidic chamber (p<0.001; n=6). Reported pairwise comparisons are the result of Wilcoxon signed-rank test. All error bars indicated are standard error.

**SI Figure S6:**
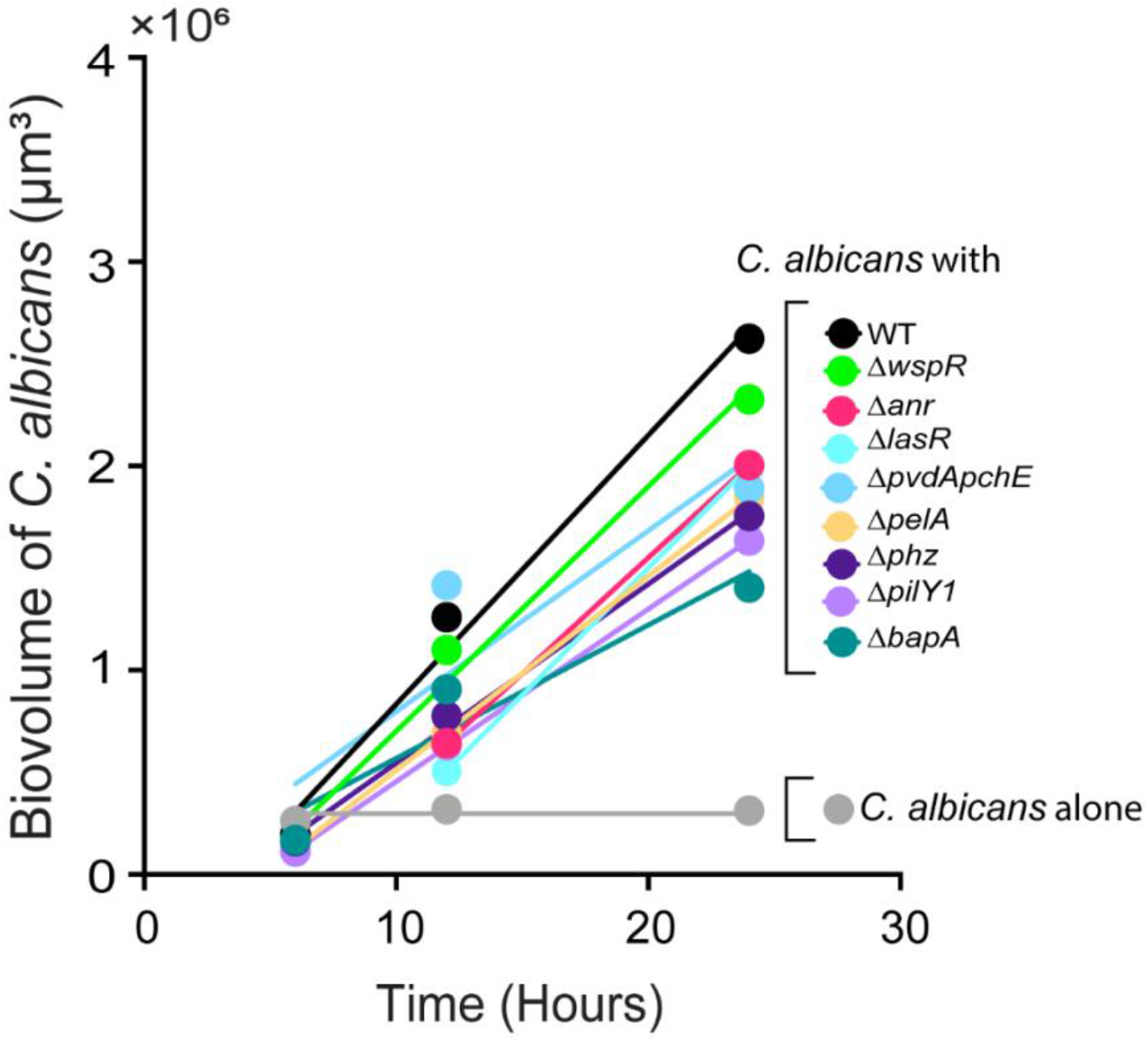
Slopes of *C. albicans* growth yields. Best fit lines were generated by fitting kinetic biofilm biovolume data by least squares regression. The slopes of the best fit lines for growth yields of *C. albicans* grown in the presence of the *P. aeruginosa* mutants were statistically different from the slope determined for *C. albicans* grown alone. (p < 0.001 for slope of *C. albicans* with *P. aeruginosa* mutants compared to slope of 0 for *C. albicans* alone as determined by extra sum-of-squares F Test).

**SI Figure S7:**
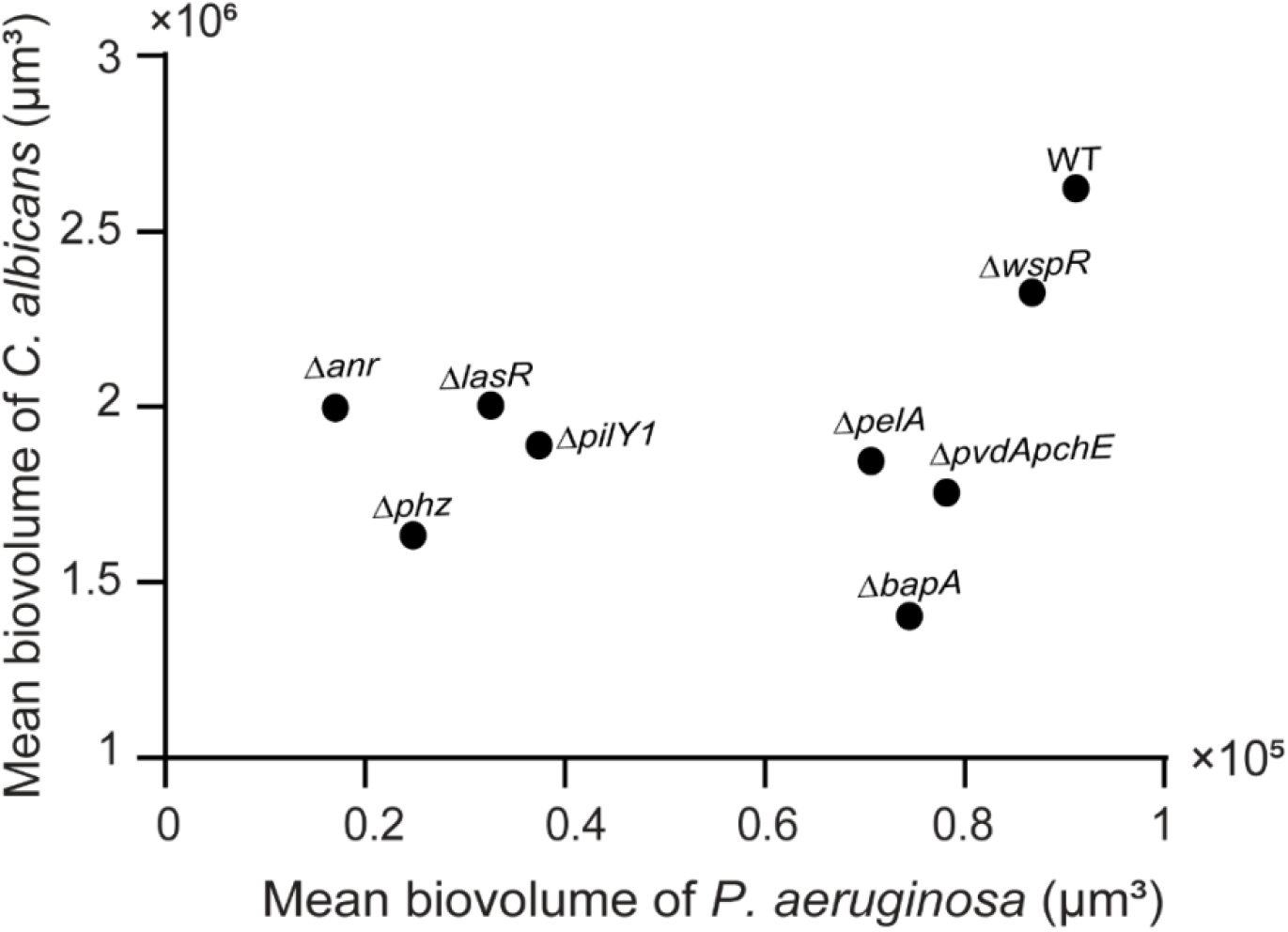
Mean *P. aeruginosa* and *C. albicans* biovolume. The mean values of biovolume of *C. albicans* mutants is plotted against the corresponding mean value of *P. aeruginosa* WT and mutant biovolume from their dual species biofilms. There is no significant correlation between the two (Linear correlation analysis; p = 0.391; r^2^= 0.107).

**SI Figure S8:**
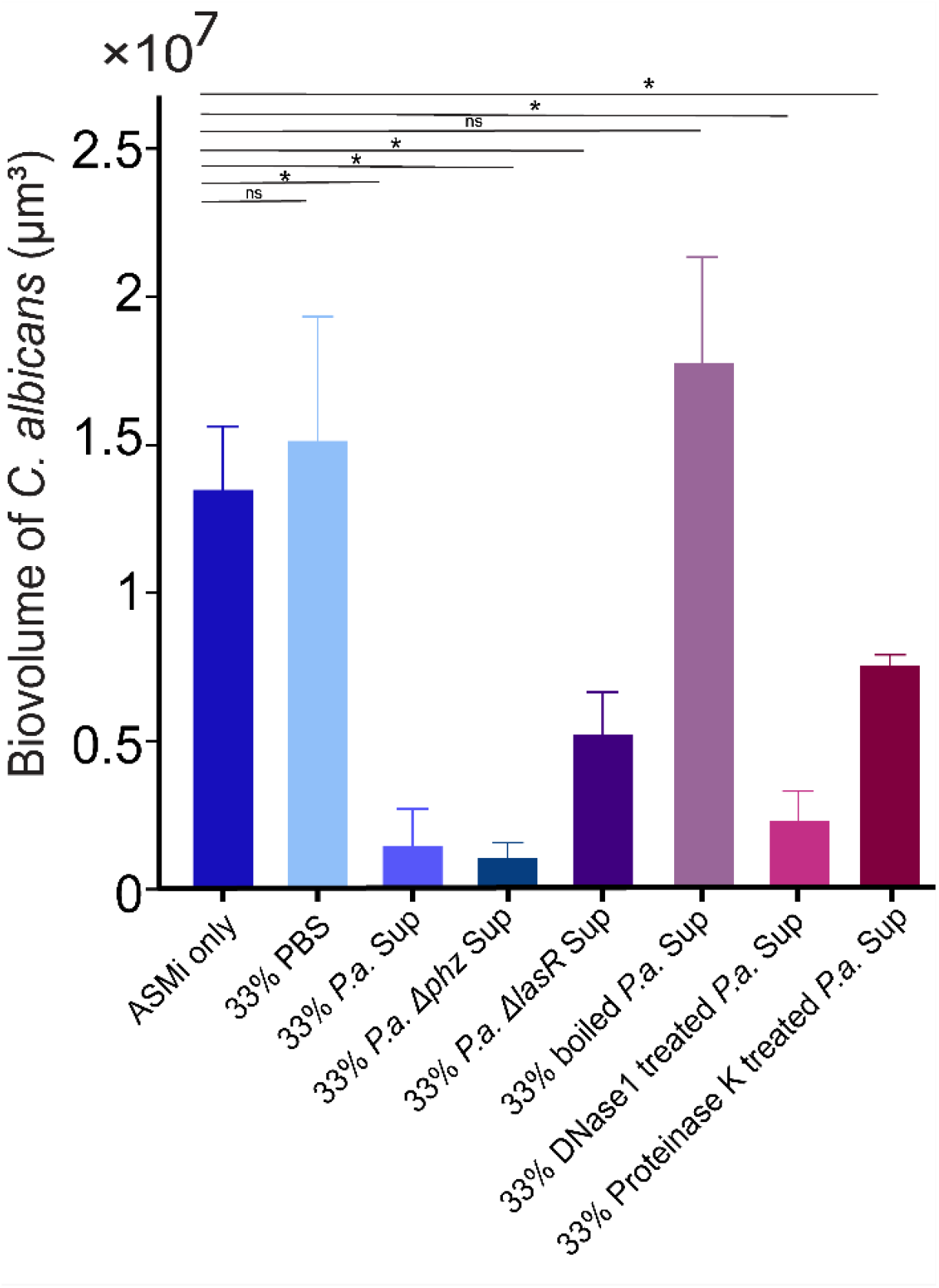
Supernatants from *P. aeruginosa* mutants and treated supernatants. Supernatants either from *P. aeruginosa* mutants or supernatant from *P. aeruginosa* PA14 wildtype treated with *Proteinase K, Dnase1* or boiled and then used to grow *C. albicans* biofilms to identify the component in the supernatant responsible for decreased *C. albicans* growth yield (* denotes p<0.05; n=6). Reported pairwise comparisons are the result of Wilcoxon signed-rank test. All error bars indicated are standard error.

**Table 1:**
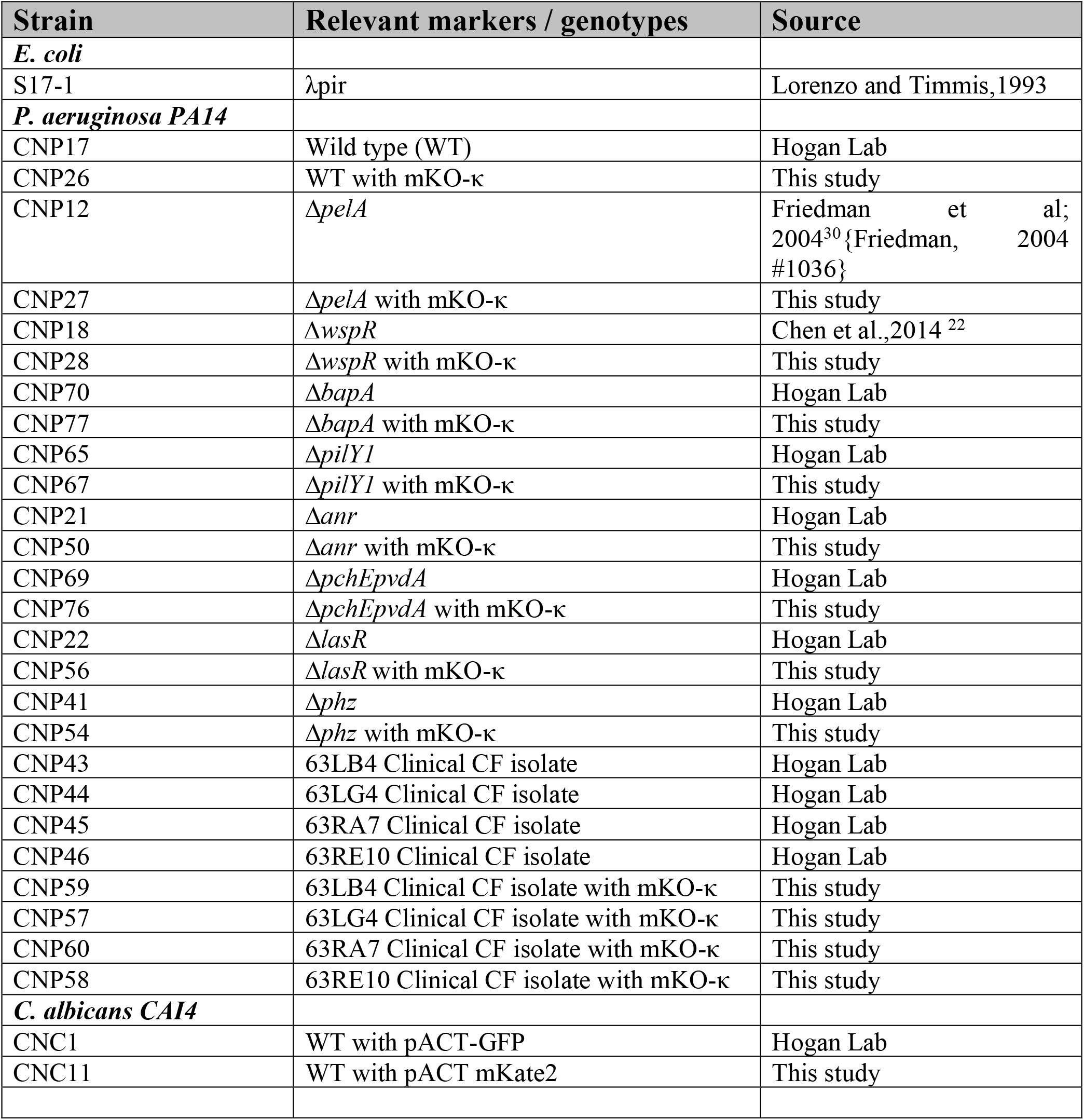
Strains and Plasmids

### Artificial Sputum media for imaging (ASMi)

#### Stocks for Base

Na_2_HPO_4_ (0.2M, 0.69g/25mL)

NaH_2_PO_4_ (0.2M, 0.71g/25mL)

KNO_3_ (1M, 2.53g/25mL)

K_2_SO_4_ (0.25M, 1.09g/25mL)

#### Additional Stocks

Glucose (20% w/v) *autoclave*

L-Lactic acid (1M) *pH to 7 with NaOH

CaCl_2_*2H_2_O (1M, 3.68g/25mL)

MgCl_2_*6H_2_O (1M, 5.08g/25mL)

FeSO_4_*7H_2_O (1mg/1mL) syringe

N-acetylglucosamine (.25M, 1.383g/25mL)

Tryptophan (0.1M, 1.021g/50mL)

#### Reagents

DNA (Herring sperm DNA)

Fucose

GalNAc

Galactose

Choline chloride

Sodium octanoate

Yeast Synthetic Dropout – Trp

NaCl

MOPS

KCl

NH 4 Cl

NaOH

#### Preparation of ASMi (500 mL) (2X in 250mL)

1. Add 400mL diH_2_O and stir bar to a clean beaker
2. While stirring add

- 3.250 mL Na_2_HPO_4_ stock
- 3.126 mL NaH2PO4 stock
- 174 uL KNO_3_ stock
- 542 uL K_2_SO_4_ stock
- 2 g Yeast Synthetic Dropout - Trp
- 1.516 g NaCl
- 1.046 g MOPS
- 558 mg KCl
- 62 mg NH_4_Cl
- 4.65 mL L-Lactic acid stock
- 1.365 mL glucose stock
- 875 uL CaCl_2_*2H_2_O stock
- 600 uL N-acetylglucosamine
- 500 uL FeSO_4_*7H_2_O
- 330 uL Tryptophan stock
- 303 uL MgCl_2_*6H_2_O
- 300mg DNA
- 0.007g Choline chloride
- 0.022g sodium octanoate
(Replacement 1,2-dipalmitoyl-sn-glycero-3-phosphocholine, DPPC)
- 400 mg Fucose
- 125 mg GalNAc
- 90 mg Galactose
(Replacement for mucin; these are mucin sugars)
3. pH to 6.8 with HCl or NaOH and add diH_2_O to 500 mL
4. Filter sterilize

#### Considerations and References

- Lacks Sphingolipids and surfactant proteins, which are moderately abundant
- Mucin sugars are used instead of mucin. [5]
- Reports of some concentrations vary from source to source

DPPC 10ug/ml [4], 100ug/ml [1]*, 2.3mg/ml [3] (Octanoate and choline are used instead at the same concentrations; 2:1:: Octanoate: Choline, since DPPC has two lipid chains per choline. DPPC molarity for Choline and 2X that for Octanoate)

DNA 600ug/ml [1]*, 2.7mg/ml [3], 4mg/ml [2]

Mucin 5mg/1ml [1]*[2] – 16mg/ml [3]

*Concentrations used in this recipe

#### Footnotes

1 http://www.pnas.org/content/suppl/2015/03/11/1419677112.DCSupplemental/pnas.1419677112.sd08.xlsx

2 http://www.nature.com/protocolexchange/protocols/1999#/procedure

3 https://www.atsjournals.org/doi/pdf/10.1164/ajrccm.164.3.2011041

4 https://www.atsjournals.org/doi/pdf/10.1164/ajrccm.164.3.2011041

5 https://www.physiology.org/doi/full/10.1152/ajplung.00108.2004?url_ver=Z39.882003&rfr_id=ori:rid:crossref.org&rfr_dat=cr)pub%3dpubmed

## References

1. Nadell, C. D., Drescher, K. & Foster, K. R. Spatial structure, cooperation and competition in biofilms. Nat. Rev. Microbiol. 14, 589–600 (2016).

2. Dufrêne, Y. F. & Persat, A. Mechanomicrobiology: how bacteria sense and respond to forces. Nat. Rev. Microbiol. (2020) doi:10.1038/s41579-019-0314-2.

3. Flemming, H.-C. et al. Biofilms: an emergent form of bacterial life. Nat. Rev. Microbiol. 14, 563–575 (2016).

4. Elias, S. & Banin, E. Multi-species biofilms: living with friendly neighbors. FEMS Microbiol. Rev. 36, 990–1004 (2012).

5. Bispo, P., Haas, W. & Gilmore, M. Biofilms in Infections of the Eye. Pathogens 4, 111–136 (2015).

6. Bowen, W. H., Burne, R. A., Wu, H. & Koo, H. Oral Biofilms: Pathogens, Matrix, and Polymicrobial Interactions in Microenvironments. Trends Microbiol. 26, 229–242 (2018).

7. Silverstein, A. & Donatucci, C. F. Bacterial Biofilms and implantable prosthetic devices. Int. J. Impot. Res. 15, S150–S154 (2003).

8. Zhao, G. et al. Biofilms and Inflammation in Chronic Wounds. Adv. Wound Care 2, 389–399 (2013).

9. Filkins, L. M. & O’Toole, G. A. Cystic Fibrosis Lung Infections: Polymicrobial, Complex, and Hard to Treat. PLOS Pathog. 11, e1005258 (2015).

10. Ciofu, O., Tolker-Nielsen, T., Jensen, P. Ø., Wang, H. & Høiby, N. Antimicrobial resistance, respiratory tract infections and role of biofilms in lung infections in cystic fibrosis patients. Adv. Drug Deliv. Rev. 85, 7–23 (2015).

11. Stacy, A., McNally, L., Darch, S. E., Brown, S. P. & Whiteley, M. The biogeography of polymicrobial infection. Nat. Rev. Microbiol. 14, 93–105 (2016).

12. Grahl, N. et al. Profiling of Bacterial and Fungal Microbial Communities in Cystic Fibrosis Sputum Using RNA. mSphere 3, e00292–18 (2018).

13. Cutting, G. R. Cystic fibrosis genetics: from molecular understanding to clinical application. Nat. Rev. Genet. 16, 45–56 (2015).

14. Peters, B. M., Jabra-Rizk, M. A., O’May, G. A., Costerton, J. W. & Shirtliff, M. E. Polymicrobial Interactions: Impact on Pathogenesis and Human Disease. Clin. Microbiol. Rev. 25, 193–213 (2012).

15. Fourie & Pohl. Beyond Antagonism: The Interaction Between Candida Species and Pseudomonas aeruginosa. J. Fungi 5, 34 (2019).

16. Pierce, G. E. Pseudomonas aeruginosa, Candida albicans, and device-related nosocomial infections: implications, trends, and potential approaches for control. J. Ind. Microbiol. Biotechnol. 32, 309–318 (2005).

17. Nobile, C. J. & Johnson, A. D. *Candida albicans* Biofilms and Human Disease. Annu. Rev. Microbiol. 69, 71–92 (2015).

18. Moradali, M. F., Ghods, S. & Rehm, B. H. A. Pseudomonas aeruginosa Lifestyle: A Paradigm for Adaptation, Survival, and Persistence. Front. Cell. Infect. Microbiol. 7, (2017).

19. Christiaen, S. E. A. et al. Bacteria that inhibit quorum sensing decrease biofilm formation and virulence in *Pseudomonas aeruginosa* PAO1. Pathog. Dis. 70, 271–279 (2014).

20. Hogan, D. A. & Kolter, R. Interactions: An Ecological Role for Virulence Factors. 296, 5 (2002).

21. Grahl, N. et al. Mitochondrial Activity and Cyr1 Are Key Regulators of Ras1 Activation of C. albicans Virulence Pathways. PLOS Pathog. 11, e1005133 (2015).

22. Chen, A. I. et al. Candida albicans Ethanol Stimulates Pseudomonas aeruginosa WspR-Controlled Biofilm Formation as Part of a Cyclic Relationship Involving Phenazines. PLoS Pathog. 10, e1004480 (2014).

23. Lewis, K. A. et al. Ethanol Decreases *Pseudomonas aeruginosa* Flagellar Motility through the Regulation of Flagellar Stators. J. Bacteriol. 201, e00285–19, /jb/201/18/JB.00285-19.atom (2019).

24. Bergeron, A. C. et al. Candida albicans and Pseudomonas aeruginosa Interact To Enhance Virulence of Mucosal Infection in Transparent Zebrafish. Infect. Immun. 85, e00475–17, /iai/85/11/e00475-17.atom (2017).

25. Turner, K. H., Wessel, A. K., Palmer, G. C., Murray, J. L. & Whiteley, M. Essential genome of *Pseudomonas aeruginosa* in cystic fibrosis sputum. Proc. Natl. Acad. Sci. 112, 4110–4115 (2015).

26. Darch, S. E. et al. Spatial determinants of quorum signaling in a *Pseudomonas aeruginosa* infection model. Proc. Natl. Acad. Sci. 115, 4779–4784 (2018).

27. Flynn, J. M., Niccum, D., Dunitz, J. M. & Hunter, R. C. Evidence and Role for Bacterial Mucin Degradation in Cystic Fibrosis Airway Disease. PLOS Pathog. 12, e1005846 (2016).

28. Hogan, D. A., Vik, Å. & Kolter, R. A Pseudomonas aeruginosa quorum-sensing molecule influences Candida albicans morphology: Pseudomonas inhibition of C. albicans filamentation. Mol. Microbiol. 54, 1212–1223 (2004).

29. Morales, D. K. & Hogan, D. A. Candida albicans Interactions with Bacteria in the Context of Human Health and Disease. PLoS Pathog. 6, e1000886 (2010).

30. Friedman, L. & Kolter, R. Genes involved in matrix formation in Pseudomonas aeruginosa PA14 biofilms. Mol. Microbiol. 51, 675–690 (2004).

31. Jennings, L. K. et al. Pel is a cationic exopolysaccharide that cross-links extracellular DNA in the *Pseudomonas aeruginosa* biofilm matrix. Proc. Natl. Acad. Sci. 112, 11353–11358 (2015).

32. Hickman, J. W., Tifrea, D. F. & Harwood, C. S. A chemosensory system that regulates biofilm formation through modulation of cyclic diguanylate levels. Proc. Natl. Acad. Sci. 102, 14422–14427 (2005).

33. Jackson, A. A. et al. Anr and Its Activation by PlcH Activity in Pseudomonas aeruginosa Host Colonization and Virulence. J. Bacteriol. 195, 3093–3104 (2013).

34. Dietrich, L. E. P. et al. Bacterial Community Morphogenesis Is Intimately Linked to the Intracellular Redox State. J. Bacteriol. 195, 1371–1380 (2013).

35. de Bentzmann, S. et al. Unique Biofilm Signature, Drug Susceptibility and Decreased Virulence in Drosophila through the Pseudomonas aeruginosa Two-Component System PprAB. PLoS Pathog. 8, e1003052 (2012).

36. Rodesney, C. A. et al. Mechanosensing of shear by *Pseudomonas aeruginosa* leads to increased levels of the cyclic-di-GMP signal initiating biofilm development. Proc. Natl. Acad. Sci. 114, 5906–5911 (2017).

37. O’Loughlin, C. T. et al. A quorum-sensing inhibitor blocks Pseudomonas aeruginosa virulence and biofilm formation. Proc. Natl. Acad. Sci. 110, 17981–17986 (2013).

38. Harrison, F. & Buckling, A. Siderophore production and biofilm formation as linked social traits. ISME J. 3, 632–634 (2009).

39. Gibson, J., Sood, A. & Hogan, D. A. Pseudomonas aeruginosa-Candida albicans Interactions: Localization and Fungal Toxicity of a Phenazine Derivative. Appl. Environ. Microbiol. 75, 504–513 (2009).

40. Morales, D. K. et al. Control of Candida albicans Metabolism and Biofilm Formation by Pseudomonas aeruginosa Phenazines. mBio 4, e00526–12 (2013).

41. Jovanovic, M. et al. Rhamnolipid inspired lipopeptides effective in preventing adhesion and biofilm formation of Candida albicans. Bioorganic Chem. 87, 209–217 (2019).

42. Hartmann, R. et al. BiofilmQ, a software tool for quantitative image analysis of microbial biofilm communities. http://biorxiv.org/lookup/doi/10.1101/735423 (2019) doi:10.1101/735423.

43. Yang, L. et al. Current understanding of multi-species biofilms. Int. J. Oral Sci. 3, 74–81 (2011).

44. Peleg, A. Y., Hogan, D. A. & Mylonakis, E. Medically important bacterial–fungal interactions. Nat. Rev. Microbiol. 8, 340–349 (2010).

45. Trejo-Hernández, A., Andrade-Domínguez, A., Hernández, M. & Encarnación, S. Interspecies competition triggers virulence and mutability in Candida albicans–Pseudomonas aeruginosa mixed biofilms. ISME J. 8, 1974–1988 (2014).

46. Nadell, C. D., Xavier, J. B. & Foster, K. R. The sociobiology of biofilms. FEMS Microbiol. Rev. 33, 206–224 (2009).

47. Oliveira, N. M. et al. Biofilm Formation As a Response to Ecological Competition. PLOS Biol. 13, e1002191 (2015).

48. Schluter, J., Nadell, C. D., Bassler, B. L. & Foster, K. R. Adhesion as a weapon in microbial competition. ISME J 9, 139–149 (2015).

49. Coyte, K. Z., Tabuteau, H., Gaffney, E. A., Foster, K. R. & Durham, W. M. Microbial competition in porous environments can select against rapid biofilm growth. Proc. Natl. Acad. Sci. (2016) doi:10.1073/pnas.1525228113.

50. Nadell, C. D., Ricaurte, D., Yan, J., Drescher, K. & Bassler, B. L. Flow environment and matrix structure interact to determine spatial competition in Pseudomonas aeruginosa biofilms. doi:10.7554/eLife.21855.001.

51. Al-Fattani, M. A. Biofilm matrix of Candida albicans and Candida tropicalis: chemical composition and role in drug resistance. J. Med. Microbiol. 55, 999–1008 (2006).

52. Zarnowski, R. et al. Candida albicans biofilm–induced vesicles confer drug resistance through matrix biogenesis. PLOS Biol. 16, e2006872 (2018).

53. Mitchell, K. F. et al. Community participation in biofilm matrix assembly and function. Proc. Natl. Acad. Sci. 112, 4092–4097 (2015).

54. Rusconi R, Stocker R. Microbes in flow. Curr. Opin. Microbiol. 25, 1–8.

55. Wheeler, J. D., Secchi, E., Rusconi, R. & Stocker, R. Not Just Going with the Flow: The Effects of Fluid Flow on Bacteria and Plankton. Annu. Rev. Cell Dev. Biol. 35, 213–237 (2019).

56. Yawata, Y., Nguyen, J., Stocker, R. & Rusconi, R. Microfluidic Studies of Biofilm Formation in Dynamic Environments. J. Bacteriol. 198, 2589–2595 (2016).

57. Rusconi, R., Garren, M. & Stocker, R. Microfluidics Expanding the Frontiers of Microbial Ecology. Annu. Rev. Biophys. 43, 65–91 (2014).

58. Persat, A. et al. The Mechanical World of Bacteria. Cell 161, 988–997 (2015).

59. Nadell, C. D. et al. Cutting through the complexity of cell collectives. Proc. R. Soc. B Biol. Sci. 280, 20122770 (2013).

60. Hartmann, R. et al. Emergence of three-dimensional order and structure in growing biofilms. Nat. Phys. 15, 251–256 (2019).

61. Liu, Y. & Tay, J.-H. The essential role of hydrodynamic shear force in the formation of biofilm and granular sludge. Water Res. 36, 1653–1665 (2002).

62. Stewart, P. S. Biophysics of biofilm infection. Pathog. Dis. 70, 212–218 (2014).

63. Ebrahimi, A., Schwartzman, J. & Cordero, O. X. Cooperation and spatial self-organization determine rate and efficiency of particulate organic matter degradation in marine bacteria. Proc. Natl. Acad. Sci. 116, 23309–23316 (2019).

64. Kannan, A., Yang, Z., Kim, M. K., Stone, H. A. & Siryaporn, A. Dynamic switching enables efficient bacterial colonization in flow. Proc. Natl. Acad. Sci. 115, 5438–5443 (2018).

65. Persat, A., Stone, H. A. & Gitai, Z. The curved shape of Caulobacter crescentus enhances surface colonization in flow. Nat. Commun. 5, 3824 (2014).

66. Secchi, E. et al. The effect of flow on swimming bacteria controls the initial colonization of curved surfaces. Nat. Commun. 11, 2851 (2020).

67. Nadell, C. D., Ricaurte, D., Yan, J., Drescher, K. & Bassler, B. L. Flow environment and matrix structure interact to determine spatial competition in Pseudomonas aeruginosa biofilms. eLife 6, e21855 (2017).

68. Martínez-García, R., Nadell, C. D., Hartmann, R., Drescher, K. & Bonachela, J. A. Cell adhesion and fluid flow jointly initiate genotype spatial distribution in biofilms. PLOS Comput. Biol. 14, e1006094 (2018).

69. Wucher, B. R. et al. Vibrio cholerae filamentation promotes chitin surface attachment at the expense of competition in biofilms. Proc. Natl. Acad. Sci. 116, 14216–14221 (2019).

70. Young, K. D. The Selective Value of Bacterial Shape. Microbiol. Mol. Biol. Rev. 70, 660–703 (2006).

71. Yang, D. C., Blair, K. M. & Salama, N. R. Staying in Shape: the Impact of Cell Shape on Bacterial Survival in Diverse Environments. Microbiol. Mol. Biol. Rev. 80, 187–203 (2016).

72. Rusconi, R., Guasto, J. S. & Stocker, R. Bacterial transport suppressed by fluid shear. Nat. Phys. 10, 212–217 (2014).

73. Besemer, K., Singer, G., Hï¿½dl, I. & Battin, T. J. Bacterial Community Composition of Stream Biofilms in Spatially Variable-Flow Environments. Appl. Environ. Microbiol. 75, 7189–7195 (2009).

74. Besemer, K. et al. Biophysical Controls on Community Succession in Stream Biofilms. Appl. Environ. Microbiol. 73, 4966–4974 (2007).

75. Rossy, T., Nadell, C. D. & Persat, A. Cellular advective-diffusion drives the emergence of bacterial surface colonization patterns and heterogeneity. Nat. Commun. 10, 2471 (2019).

76. Besemer, K. et al. Unraveling assembly of stream biofilm communities. ISME J. 6, 1459–1468 (2012).

77. Lecuyer S, Rusconi R, Shen Y, Forsyth A, Vlamakis H, Kolter R, Stone HA. Shear Stress Increases the Residence Time of Adhesion of Pseudomonas aeruginosa. Biophys. J. 100, 341–350.

78. Siryaporn, A., Kim, M. K., Shen, Y., Stone, H. A. & Gitai, Z. Colonization, competition, and dispersal of pathogens in fluid flow networks. Curr. Biol. 25, 1201–1207 (2015).

79. Dingemans, J. et al. Effect of Shear Stress on *Pseudomonas aeruginosa* Isolated from the Cystic Fibrosis Lung. mBio 7, e00813–16, /mbio/7/4/e00813-16.atom (2016).

80. Mukherjee, P. K., Chand, D. V., Chandra, J., Anderson, J. M. & Ghannoum, M. A. Shear stress modulates the thickness and architecture of Candida albicans biofilms in a phase-dependent manner. Mycoses 52, 440–446 (2009).

81. Alves, P. M. et al. Interaction between Staphylococcus aureus and Pseudomonas aeruginosa is beneficial for colonisation and pathogenicity in a mixed biofilm. Pathog. Dis. 76, (2018).

82. Carolus, H., Van Dyck, K. & Van Dijck, P. Candida albicans and Staphylococcus Species: A Threatening Twosome. Front. Microbiol. 10, 2162 (2019).

83. Orazi, G., Ruoff, K. L. & O’Toole, G. A. *Pseudomonas aeruginosa* Increases the Sensitivity of Biofilm-Grown *Staphylococcus aureus* to Membrane-Targeting Antiseptics and Antibiotics. mBio 10, e01501–19, /mbio/ 10/4/mBio.01501-19.atom (2019).

84. Shing, S. R. et al. The Fungal Pathogen Candida albicans Promotes Bladder Colonization of Group B Streptococcus. Front. Cell. Infect. Microbiol. 9, 437 (2020).

85. Scoffield, J. A., Duan, D., Zhu, F. & Wu, H. A commensal streptococcus hijacks a Pseudomonas aeruginosa exopolysaccharide to promote biofilm formation. PLOS Pathog. 13, e1006300 (2017).

86. Azoulay, E. et al. Candida Colonization of the Respiratory Tract and Subsequent Pseudomonas Ventilator-Associated Pneumonia. Chest 129, 110–117 (2006).

87. Earle, K. A. et al. Quantitative Imaging of Gut Microbiota Spatial Organization. Cell Host Microbe 18, 478–488 (2015).

88. Welch, J. L. M., Rossetti, B. J., Rieken, C. W., Dewhirst, F. E. & Borisy, G. G. Biogeography of a human oral microbiome at the micron scale. Proc. Natl. Acad. Sci. 113, E791–E800 (2016).

89. Welch, J. L. M., Hasegawa, Y., McNulty, N. P., Gordon, J. I. & Borisy, G. G. Spatial organization of a model 15-member human gut microbiota established in gnotobiotic mice. Proc. Natl. Acad. Sci. 114, E9105–E9114 (2017).

90. Gallego-Hernandez, A. L. et al. Upregulation of virulence genes promotes Vibrio cholerae biofilm hyperinfectivity. Proc. Natl. Acad. Sci. 117, 11010–11017 (2020).

91. Caldara, M. et al. Mucin Biopolymers Prevent Bacterial Aggregation by Retaining Cells in the Free-Swimming State. Curr. Biol. 22, 2325–2330 (2012).

92. Wheeler, K. M. et al. Mucin glycans attenuate the virulence of Pseudomonas aeruginosa in infection. Nat. Microbiol. 4, 2146–2154 (2019).

93. Shanks, R. M. Q., Caiazza, N. C., Hinsa, S. M., Toutain, C. M. & O’Toole, G. A. Saccharomyces cerevisiae-Based Molecular Tool Kit for Manipulation of Genes from Gram-Negative Bacteria. Appl. Environ. Microbiol. 72, 5027–5036 (2006).

94. Barelle, C. J. et al. GFP as a quantitative reporter of gene regulation inCandida albicans. Yeast 21, 333–340 (2004).

95. Backer, M. D. D., Maes, D., Vandoninck, S., Logghe, M. & Contreras, R. Transformation of Candida albicans by electroporation. 10 (1999).

96. Ng, J. M. K., Gitlin, I., Stroock, A. D. & Whitesides, G. M. Components for integrated poly(dimethylsiloxane) microfluidic systems. ELECTROPHORESIS 23, 3461–3473 (2002).

97. O’Toole, G. A. Microtiter Dish Biofilm Formation Assay. J. Vis. Exp. 2437 (2011) doi: 10.3791/2437.

